# STITCH: Spatial Transcriptomics Imputation via Flow Matching with Internal Learning

**DOI:** 10.64898/2026.06.03.729557

**Authors:** Sui Wang, Xinyu Wang, Qiangwei Peng, Tiejun Li

## Abstract

Spatial transcriptomics datasets frequently suffer from spatial gaps and missing regions due to sectioning artifacts, tissue damage, and the high cost of sequencing that limits tissue coverage. We present STITCH, a scalable and robust generative framework for multidimensional virtual spatial transcriptomics reconstruction. STITCH models intrinsic spatial-transcriptomic patterns directly from individual tissue samples, enabling reconstruction without requiring external reference atlases or matched histological image priors. The framework adopts a decoupled architecture that separates spatial morphology restoration from transcriptomic generation. STITCH first compresses high dimensional transcriptomic profiles into a low-dimensional latent representation through a spatial-aware graph autoencoder. For 3D cross-slice gaps, STITCH employs optimal transport-conditioned flow matching for spatial reconstruction, whereas 2D in-slice damage is repaired through an internal learning strategy. To generate the corresponding transcriptomic profiles, STITCH further establishes a point-wise conditional flow matching model in the latent space. This module achieves linear computational complexity, enabling continuous 3D atlas reconstruction of over 11 million cells within 5 hours on a single commodity GPU. Extensive evaluations across diverse spatial transcriptomics platforms, spanning both single-cell and spot-level technologies, demonstrate that STITCH consistently preserves transcriptomic identities, spatial topologies, and anatomical continuity. Overall, STITCH provides a scalable and platform-compatible computational framework for reconstructing high-resolution continuous spatial transcriptomic atlases.

## 1 Introduction

Spatial transcriptomics (ST) technologies have resolved tissue architectures at unprecedented resolutions. However, due to the constraints of physical sectioning techniques and prohibitive sequencing costs, the construction of complete and continuous macroscopic three-dimensional (3D) atlases remains severely hindered by massive “data *lacuna*”[1, 2, 3, 4, 5]. Researchers are typically limited to sparse two-dimensional (2D) slice sampling of biological tissues. Even within a single slice, technical artifacts often lead to low-quality physical voids. This sampling paradigm inevitably introduces substantial information gaps across multidimensional physical space, resulting in fragmented and highly discontinuous reconstructed tissue architectures. This severely impedes the in-depth exploration of continuous spatial manifolds and large-scale biological structures.

To address these spatial gaps, early computational methods primarily focused on slice alignment, while more recent approaches increasingly leverage high-resolution histological images for cross-modal mapping. Pioneering algorithms such as PASTE [6], STAligner [7], STitch3D[8], STalign [9], Spateo [10], and STAIR [11] have been developed based on various mathematical frameworks, including optimal transport, graph neural networks, and diffeomorphic mapping, to achieve alignment between slices or samples from different time points. However, such alignments inherently emphasize batch correction and physical superimposition between samples, leaving depth gaps of tens to hundreds of micrometers between slices. To further improve spatial continuity and resolution, a series of cross-modal mapping frameworks have been developed. Advanced frameworks such as ST-Net [12], HisToGene [13], Hist2ST [14], PRECAST [15], THItoGene [16], iStar [17], iScale [18], GHIST [19], and STimage [20] have successfully leveraged high-resolution histological images (e.g., H&E staining) as priors to achieve leaps in spatial resolution, or even directly predict high-fidelity spatial gene expression from standard images.

In many cases, we lack matched histological images as prior information, yet spatial reconstruction remains crucial. This situation is common in spatiotemporal dynamic inference tasks, which rely on high-quality ST data at observed time points [21, 22, 23]. Under such constraints, we must depend solely on the discrete spatial transcriptomic data itself and perform reconstruction directly from the intrinsic spatial context.

Several recent generative frameworks have been proposed to address this issue. For example, SpatialZ [24] combines optimal transport (OT)-based physical coordinate interpolation with probabilistic sampling that leverages local microenvironmental similarities to generate virtual slices. isoST [25] employs stochastic differential equations to model gene expression continuity across depths, while STADiffuser [26] introduces latent diffusion models to simulate 3D spatial data. Despite notable progress, purely probabilistic interpolation and multi-step diffusion generation often struggle to accurately model complex nonlinear morphological evolution and may introduce structural artifacts during cross-slice reconstruction. Moreover, the diffusion-based or SDE-based models (e.g., STADiffuser and isoST) often incur prohibitive computational and memory costs when scaled to large spatial atlases. Additionally, existing models tend to focus on single-dimensional absences, lacking a unified architecture capable of flexibly addressing complex defects, such as 3D cross-slice gaps or 2D single-slice damage.

Here, we present STITCH (Spatial Transcriptomics Imputation via flow maTCHing), a scalable and robust reconstruction framework for multidimensional virtual spatial transcriptomics generation. Our approach is inspired by internal learning methods from computer vision, such as SinGAN [27] and SinDiffusion [28], which adopt a single-sample internal learning strategy for image generations. We leverage the fact that spatial continuity, coherent microenvironmental composition, and region-specific expression architectures collectively provide sufficient internal statistical structure to support conditional generative reconstruction directly from a single observed tissue sample. This motivates our method to reconstruct missing regions by harnessing the spatial and transcriptomic coherence inherent in the observed tissue context, operating entirely within an internal learning paradigm.

STITCH adopts a decoupled architecture that separates spatial structure reconstruction from gene feature generation. First, it compresses high-dimensional gene expression profiles into a continuous low-dimensional latent space through a spatial-aware graph autoencoder. Subsequently, during the spatial structure reconstruction stage, STITCH flexibly handles different types of physical defects. For 3D cross-slice morphological evolution, STITCH employs optimal transport-conditioned flow matching (OT-CFM) [29, 30] to establish a *Structure Flow Module* for spatial coordinate inference across slices. For 2D in-slice damage, it adopts an internal learning strategy [27, 28, 31] adapted from computer vision for spatial coordinate restoration. To improve scalability for large-scale atlas reconstruction, STITCH further establishes a *Gene Flow Module* based on point-wise conditional flow matching as the core transcriptomic generative engine. Owing to this decoupled point-wise formulation, STITCH achieves linear computational complexity, enabling efficient scaling to large spatial atlases on a single commodity GPU. In addition, we construct an exploratory neighborhood-enhanced variant (STITCH-n) to evaluate the influence of local spatial neighborhood on generative performance.

We evaluate STITCH across multiple cutting-edge spatial transcriptomics platforms, including single-cell resolution technologies (Stereo-seq on *Drosophila* embryo, MERFISH and Xenium on mouse brain) and spot-level profiling datasets (Visium on human breast cancer and on human DLPFC). Across these diverse platforms, STITCH consistently preserves transcriptomic identities, spatial topologies, and anatomical continuity. Notably, by expanding a 54-slice MERFISH mouse brain dataset (containing 1,056,520 cells) into a continuous atlas of 571 slices (comprising 11,549,855 cells) within approximately 5.09 hours, STITCH demonstrates its ability to reconstruct large-scale continuous tissue structures from sparse spatial observations. Overall, STITCH provides a scalable and platform-compatible computational framework for continuous spatial transcriptomic atlas reconstruction.

## 2 Results

### 2.1 Overview of STITCH Workflow

The aim of STITCH is to present a scalable and robust generative framework for the computational reconstruction of continuous spatial tissues from sparse and incomplete observations. In general, spatial transcriptomic sequencing data suffer from data loss due to sparse slice sampling and physical damage, manifesting as wide inter-slice gaps in 3D tissue sampling and irregular intra-slice structural damage arising from experimental artifacts. STITCH resolves these issues by leveraging local geometric and transcriptomic continuity in neighboring tissue regions, which can be learned and generalized from the available samples through an internal learning paradigm. Importantly, STITCH is highly versatile and applicable to diverse sequencing modalities, spanning both single-cell and spot-level spatial transcriptomics platforms — including MERFISH [2], Stereo-seq [3], Xenium [32], and Visium [33] (Fig. 1a).

**Figure 1.**
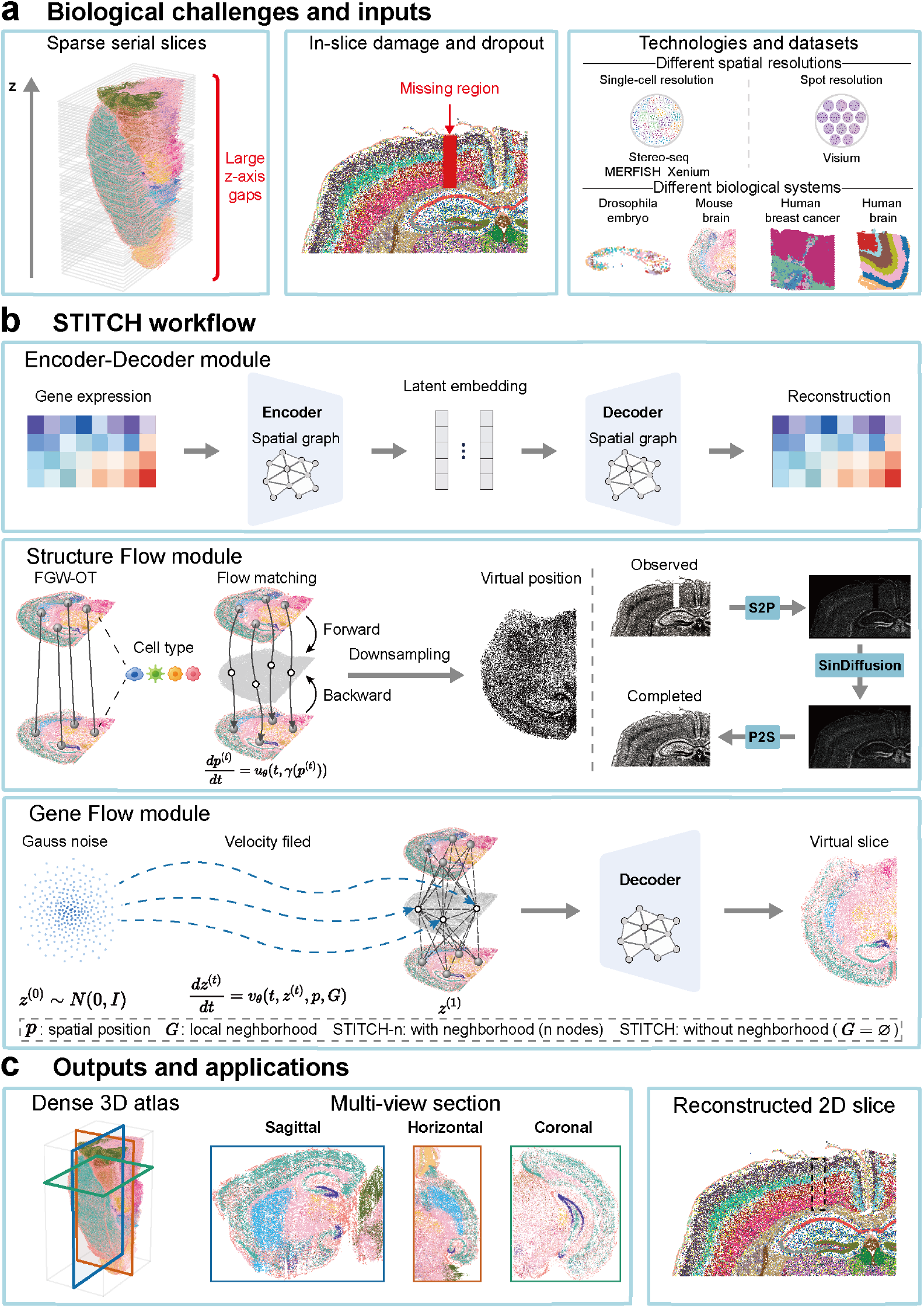
Overview of STITCH workflow. a, Biological challenges and inputs. Real-world spatial transcripttomics is limited by multi-dimensional spatial “inter-slice gaps”. The inputs include sparse 3D spatial transcriptomics data with wide inter-slice spatial gaps (left), and 2D spatial tissue slices suffering from intra-slice structural damage or experimental tears (middle). STITCH is a cross-platform framework compatible with diverse spatial transcripttomics technologies, encompassing both spot-level and single-cell resolution platforms (right). **b, The decoupled STITCH workflow**. The framework performs spatial transcriptomics reconstruction through a three-stage pipeline. **Top (Encoder-Decoder module):** Compresses high-dimensional gene expressions into a low-dimensional latent representation using a spatial-aware graph autoencoder while preserving local spatial topologies. **Middle (Structure Flow module):** Adaptively reconstructs missing spatial geometries, utilizing optimal transport flow matching for 3D cross-slice reconstruction and SinDiffusion-based reconstruction for 2D in-plane repair. **Bottom (Gene Flow module):** Establishes a continuous conditional flow matching model in the latent space to generate transcriptomic profiles conditioned on the newly generated spatial coordinates. **c, Generative outputs**. STITCH reconstructs missing spatial regions to output continuous spatial topologies. Outputs include a high-resolution continuous 3D spatial atlas (left) accompanied by continuous digital cross-sections across different orthogonal planes (middle), and a repaired 2D spatial slice recovering high-fidelity spatial coordinates and biological identities (right).

To address the inter-slice gaps and in-slice damage described above, STITCH adopts a three-stage decoupled architectECture that explicitly separates spatial morphology restoration from gene expression generation (Fig. 1b). First, the model compresses high-dimensional transcriptomic profiles into a continuous low-dimensional latent space while preserving physical topologies. To achieve this, STITCH employs a spatial-aware graph autoencoder (SAGA), thereby bypassing the computational bottlenecks associated with high-dimensional modeling (see the Encoder-Decoder module in the top panel of Fig. 1b). Second, the model restores physical coordinates according to different defect types during the spatial structure reconstruction stage. Specifically, optimal transport-conditioned flow matching is employed to reconstruct 3D cross-slice gaps, whereas an internal learning strategy is used to repair 2D in-slice damage (see the Structure Flow module in the middle panel of Fig. 1b), where “S2P” (slice-to-picture) denotes the rasterization of slice coordinates into density maps, “SinDiffusion” denotes the utilized repairing method [28], and “P2S” (picture-to-slice) denotes the decoding of reconstructed density maps back into slice coordinates. Third, conditioned on the newly inferred spatial geometry, the core Gene Flow module reconstructs the corresponding gene expression in the latent space. To this end, it leverages a scalable point-wise conditional flow matching model as its generative engine (see the Gene Flow module in the bottom panel of Fig. 1b).

Ultimately, STITCH outputs high-resolution, continuous 3D spatial atlases from which researchers can directly observe coherent digital cross-sections along arbitrary planar or curved surfaces. Alongside these 3D capabilities, STITCH also generates highly topologically restored 2D tissue slices, establishing a universal computational foundation for subsequent tissue reconstruction and downstream analysis (Fig. 1c).

### 2.2 Neighborhood-aware refinement improves generative fidelity but compromises computational efficiency

To systematically evaluate the contribution of local spatial neighborhoods to generative dynamics, we conducted a comprehensive comparative analysis between the core point-wise engine (STITCH) and its neighborhood-enhanced variant (STITCH-n) using the 3D Stereo-seq *Drosophila* embryo dataset [10], which features complex anatomical structures. This dataset comprises a total of 21,473 cells distributed across 20 slices (averaging 1,074 cells per slice). As illustrated in the ground-truth 3D atlas, real-world continuous sampling often encounters low-quality or damaged sections (e.g., slice S18), underscoring the urgent need for robust virtual reconstruction (Fig. 2a). To rigorously simulate physical inter-slice gaps, we designed a multi-level leave-out cross-validation task by progressively masking continuous slices. The gap size varied from 1 to 4 (representing the removal of slices S12, S11-S12, S10-S12, and S10-S13, respectively), imposing increasingly severe challenges on the generative models.

**Figure 2.**
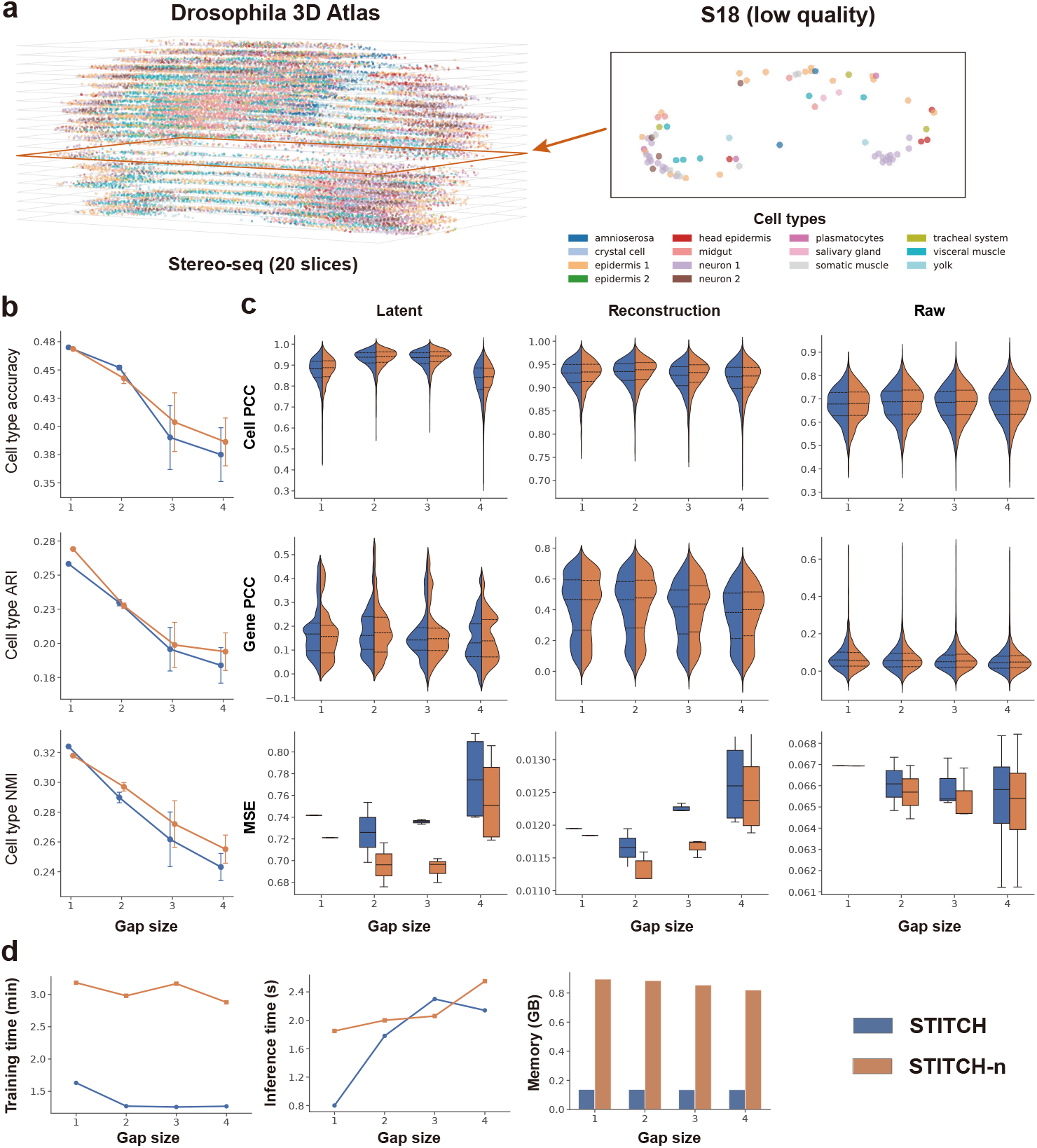
Neighborhood-aware refinement improves generative fidelity but compromises computational efficiency. **a**, The ground truth 3D spatial organization of the Stereo-seq *Drosophila* embryo (comprising 21,473 cells across 20 slices). The cell type annotations of a representative low-quality slice (S18) are highlighted, illustrating inherent physical damages during experimental acquisition. **b**, Line charts comparing the biological identity presservation of STITCH (point-wise flow) and STITCH-n (neighborhood-enhanced flow) across four progressive gap sizes (masking 1 to 4 consecutive slices). Metrics include cell type ACC, ARI, and NMI. **c**, Comprehensive evaluation of transcriptomic fidelity. A 3 *×* 3 grid displays split-violin plots (first two rows) and box plots (bottom row) comparing cell PCC, gene PCC, and MSE. The evaluations systematically span the compressed latent space, the graph autoencoder-reconstructed expression space, and the raw gene space. In the split-violin plots, the left and right halves represent STITCH and STITCH-n, respectively, with three dashed lines indicating the 25th, 50th, and 75th percentiles of the distributions. **d**, Computational profiling under identical training settings. Left: Training time overhead across different gap sizes. Middle: Inference time for reconstructing target slices (averaging ~1,000 cells per slice). Right: Peak memory consumption (GB) during generation, demonstrating the memory efficiency of STITCH.

When assessing the preservation of discrete biological identities, the neighborhood-enhanced STITCH-n demonstrated stronger resilience against wide physical faults. Both STITCH and STITCH-n maintained robust performance across the four gap sizes. Notably, while STITCH achieved a slightly higher Accuracy (ACC) at gap size 2 (0.452 vs. 0.442), STITCH-n demonstrated stronger resilience as the physical fault widened, surpassing STITCH at gap sizes 3 and 4 (0.403 vs. 0.390, and 0.386 vs. 0.375, respectively) (Fig. 2b). Similar trends in Adjusted Rand Index (ARI) and Normalized Mutual Information (NMI) confirmed that explicitly incorporating local graph topologies aids in rectifying cell identities across large missing distances.

In terms of recovering continuous transcriptomic features, the introduction of local attention enabled STITCH-n to deliver consistently higher-fidelity spatial gene expression reconstructions. To quantify this fidelity, we evaluated cell Pearson correlation coefficient (PCC), gene PCC, and mean squared error (MSE) across three distinct data space: the compressed latent space, the graph autoencoder-reconstructed expression space, and the raw gene space (Fig. 2c). In spatial omics, accurately recovering the spatial expression patterns of individual genes (gene PCC) is paramount for downstream biological discoveries. Taking gene PCC in the graph autoencoder-reconstructed expression space as a representative example, the split-violin plots reveal that STITCH-n generally elevates the overall distribution of correlations. Specifically, in most comparative scenarios, STITCH-n outperformed STITCH across the three quartile dashed lines (the 25th, 50th, and 75th percentiles). For instance, at gap size 2, the 25th, 50th, and 75th percentiles of gene PCC for STITCH-n reached 0.604, 0.484, and 0.281, respectively, demonstrating a distinct advantage over the 0.594, 0.471, and 0.280 achieved by STITCH. Similar performance improvements were also observed in the majority of evaluations for cell PCC and MSE. Overall, augmented by local attention, STITCH-n delivered a slightly refined transcriptomic reconstruction across most feature spaces.

However, incorporating local neighborhood interactions substantially increases computational overhead. Under identical hardware blocks, batch sizes, and epochs, the integration of graph attention in STITCH-n resulted in an average training time of approximately 3 minutes (Fig. 2d). In stark contrast, the decoupled architecture of STITCH drastically reduced training time to 1.2~1.65 minutes. Given that a single target slice only requires inferring roughly 1,074 cells, the absolute inference times for both models were extremely brief (ranging from 0.8 to 2.5 seconds). Notably, the graph-free STITCH achieved a fast inference time of merely 0.8 seconds at gap size 1. Crucially, STITCH exhibited an absolute scalability advantage in memory overhead: while STITCH-n consumed up to 0.8~0.9 GB of peak memory, STITCH stably required only ~0.135 GB, representing an impressive 85% reduction.

Taken together, these results establish a critical trade-off: while the local attention in STITCH-n fine-tunes generation accuracy, the point-wise STITCH core achieves highly competitive fidelity at a fraction of the computational and memory costs. Given its exceptional computational efficiency and scalability, we deploy the graph-free STITCH as the default generative engine for all subsequent multi-platform benchmarking tasks, across diverse datasets, including million-cell spatial atlases.

### 2.3 STITCH achieves accurate and scalable 3D spatial reconstruction of Drosophila dataset

To assess STITCH’s performance, we benchmarked it against SpatialZ [24] on the Drosophila dataset [10]. STITCH does not assume that slice progression strictly follows the *z*-axis, allowing accurate reconstruction even when the aligned tissue structure is spatially tilted. Therefore, to ensure a fair comparison and accommodate the algorithmic constraints of existing methods, we conducted the comparative benchmarking in Figure 3 using a standardized *z*-aligned coordinate framework. We evaluated STITCH and SpatialZ in the raw gene space across four progressive gap sizes (from 1 to 4).

**Figure 3.**
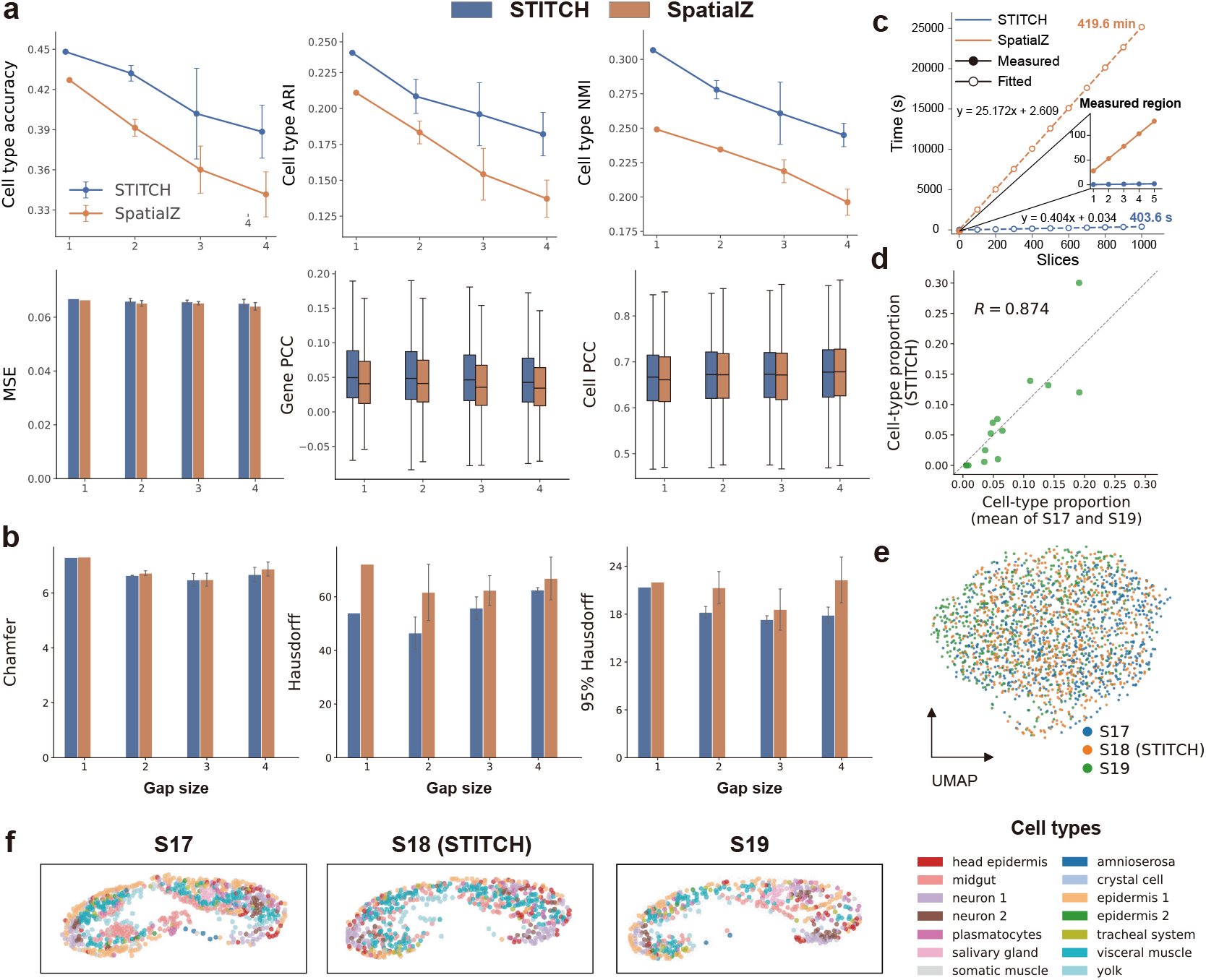
STITCH achieves accurate and scalable 3D spatial reconstruction of the Drosophila dataset. **a**, Line charts (top) and box/bar plots (bottom) comparing biological identity and transcriptomic fidelity between STITCH and SpatialZ across four progressive gap sizes (evaluated in the raw gene space). Metrics include cell type ACC, ARI, NMI, MSE, gene PCC, and cell PCC. In the box plots, the center line indicates the median. **b**, Evaluation of the generated virtual spatial structures using Chamfer distance (CD), Hausdorff distance (HD), and 95th percentile Hausdorff distance (HD95) across the four gap sizes, demonstrating STITCH’s superiority in preserving spatial geometries **c**, Extrapolation of inference time for scaling up virtual slice generation. The time required to generate 1 to 5 slices was linearly extrapolated to 1,000 slices, comparing SpatialZ with STITCH’s full end-to-end inference pipeline (Structure Flow, single-pass Gene Flow, and SAGA decoding). **d**, Scatter plot correlating the cell type proportions of the STITCH-generated virtual S18 with the contextual mean of the true S17 and S19 slices (PCC= 0.874). **e**, UMAP projection of the latent representations, illustrating that the generated S18 (orange) bridges the latent manifolds of S17 (blue) and S19 (green). **f**, Spatial visualization of key anatomical domains (e.g., midgut, neuron1, neuron2, and visceral muscle) across the reference (S17, S19) and virtual (S18) slices, showcasing continuous and biologically consistent morphological transitions.

In terms of preserving biological identity, STITCH demonstrated significant advantages (Fig. 3a, top row). Across all gap sizes, STITCH outperformed SpatialZ in ACC, ARI, and NMI. For instance, as the gap size expanded from 1 to 4, the ACC of STITCH decreased from 0.449 to 0.390, whereas SpatialZ decreased from 0.428 to 0.340. This indicates that flow matching-based generative dynamics exhibit stronger robustness when inferring cell identities across wide physical faults.

When evaluating continuous transcriptomic fidelity at the gene level (Fig. 3a, bottom row), STITCH consistently maintained a distinct advantage in capturing spatial gene distribution patterns, even though SpatialZ exhibited highly comparable or marginally lower MSE in this specific dataset. Specifically, STITCH surpassed SpatialZ in the overall distribution of gene PCC across all four gap sizes. Additionally, STITCH maintained a lead in cell PCC for the first three gap scenarios.

Regarding the morphological accuracy of the generated virtual physical coordinates, STITCH substantially outperformed SpatialZ across three rigorous geometric metrics: Chamfer distance (CD), Hausdorff distance (HD), and 95th percentile Hausdorff distance (HD95) (Fig. 3b). Notably, STITCH significantly reduced the maximum boundary error measured by HD (e.g., 53 vs. 72 at gap size 1), demonstrating that our 3D Structure Flow module exceptionally captures the non-linear morphological variations of tissue contours.

Beyond generation accuracy, computational speed is the ultimate bottleneck for transitioning to large-scale spatial atlases (Fig. 3c). We tracked the inference time required to generate 1 to 5 virtual slices and extrapolated these linear trends to reconstructing 1,000 virtual slices. The reported STITCH inference time includes all reconstruction stages, including 3D Structure Flow, Gene Flow generation, and SAGA decoding. While SpatialZ would require approximately 419.6 minutes to extrapolate 1,000 slices, the deep-learning-driven STITCH, empowered by an efficient ODE solver, completes the task in roughly 403.6 seconds.

Finally, a single-tier biological validation on the virtual slice (S18) bridging S17 and S19 confirmed that STITCH successfully recovers coherent tissue architecture. Quantitative analysis showed that the cell type proportions in the STITCH-generated S18 highly correlated with the true contextual mean of S17 and S19 (PCC=0.874) (Fig. 3d). In the latent UMAP [34] projection, the virtual S18 cells embedded themselves between the latent manifolds of S17 and S19, forging a smooth transcriptomic trajectory (Fig. 3e). Spatially, key anatomical domains, such as the midgut, neurons (neuron1 and neuron2), and visceral muscle, exhibited smooth topological transitions across the slices (Fig. 3f), vividly demonstrating STITCH’s capability to preserve topological integrity across reconstructed slices.

### 2.4 STITCH enables efficient construction of a continuous 10-million-cell scale 3D mouse brain atlas

To evaluate the scalability of STITCH and its ability to reconstruct spatial transcriptomic data with curved slice geometries, we applied it to a filtered subset of the MERFISH mouse brain dataset (Zhuang-ABCA-2 [4]) retaining 54 high-quality sections comprising 1,056,520 cells (Fig. 4a). Importantly, because this dataset were registered into the Allen Mouse Brain Common Coordinate Framework v3 (CCFv3) [35], the originally slices are represented as oblique and locally deformed sheets in CCF space rather than perfectly planar slices. Furthermore, after the removal of specific slices, spatial gaps of varying widths remained within the dataset (Fig. 4b). Addressing this real-world biological scenario, we configured STITCH to generate 9 virtual slices within each standard gap (approximately 0.2 mm) and 19 virtual slices within each of the 4 wider gaps caused by slice exclusion. To enable finer-grained generation of a specific anatomical structure of interest, such as the hippocampus, whose associated cells account for only a small fraction of the whole-brain dataset, we adopted an importance-sampling strategy. Instead of uniform spatial sampling, we assigned a higher training fraction to hippocampus-associated cells while maintaining whole-brain coverage. This strategy improved the preservation of fine hippocampal architecture, particularly within the hippocampal CA fields (CA1, CA2, CA3) and the dentate gyrus (DG) (see the corresponding uniform-sampling ablation in Supplementary Fig. S1a,b). Ultimately, STITCH successfully generated 517 virtual slices and 10,493,335 virtual cells, expanding the original data by approximately 9.93-fold. This resulted in an high-resolution 3D continuous mouse brain atlas comprising 571 slices and a total of 11,549,855 cells, with the entire 10-million-cell scale reconstruction completing in just 5.09 hours (18,340 seconds) (Fig. 4c). Representative generated results from multiple slice intervals in the whole-brain reconstruction, including both standard gaps and wider gaps caused by slice exclusion, are shown in Supplementary Fig. S1c.

**Figure 4.**
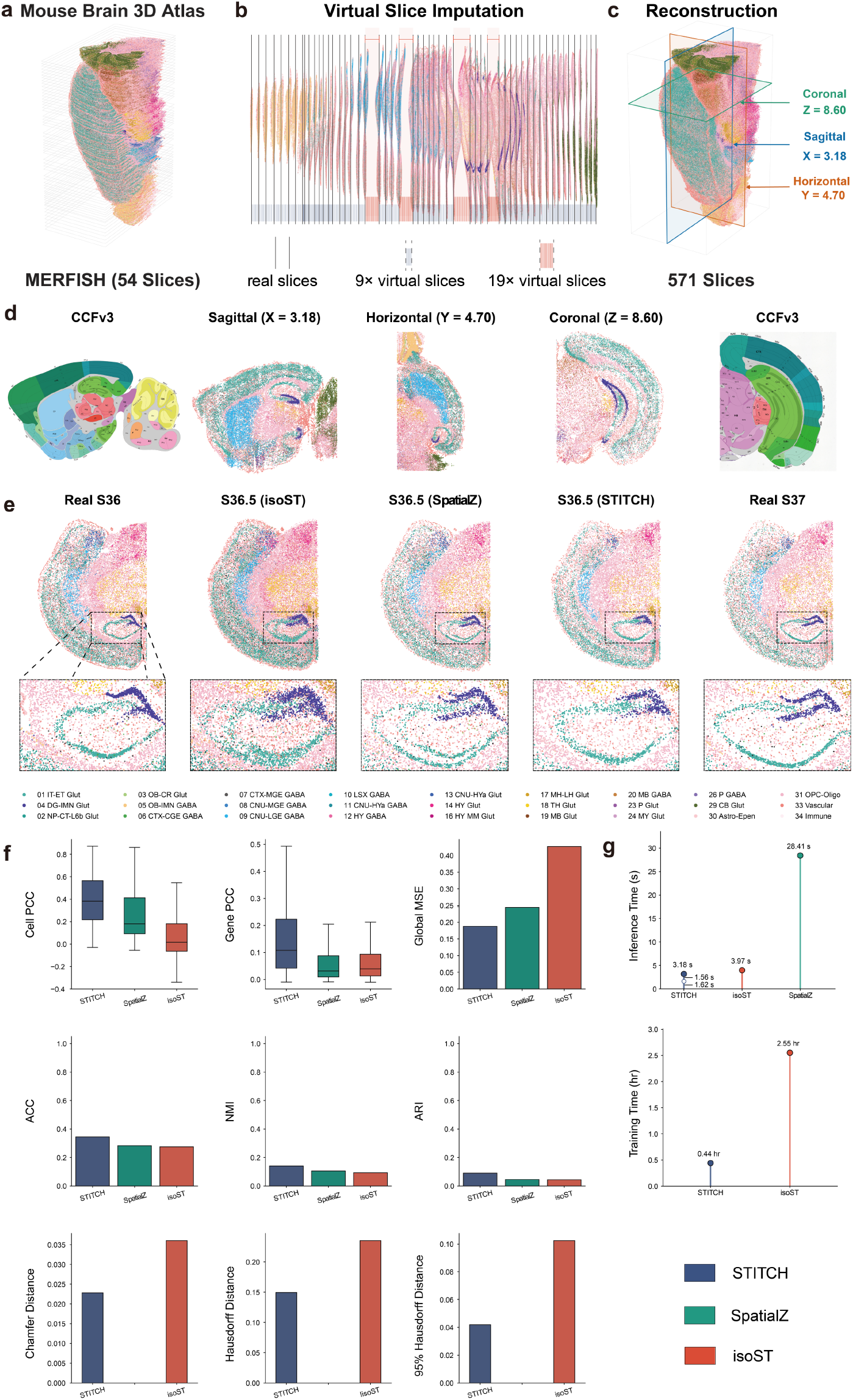
STITCH enables efficient construction of a continuous 10-million-cell scale 3D mouse brain atlas. **a**, 3D overall visualization of the raw MERFISH mouse brain (ABCA-2) dataset, comprising 54 rigorously filtered authentic slices (totaling 1,056,520 cells). **b**, Visualization of spatial gaps in the original spatial space. After registration to the Allen Mouse Brain CCFv3, the originally slices are represented as oblique and locally deformed sheets in the CCF-registered coordinate space. STITCH was configured to generate 9 virtual slices within the standard gaps and 19 virtual slices within the 4 wider gaps caused by slice exclusion. **c**, The high-resolution 3D continuous atlas reconstructed by STITCH. During whole-brain Gene Flow training, hippocampus-associated cells were assigned a higher training fraction to preserve fine hippocampal architecture while maintaining whole-brain sampling coverage. The model generated a total of 517 virtual slices, successfully expanding the dataset to 571 slices and 11,549,855 cells, all within 5.09 hours. **d**, Digital sections of the 3D atlas across orthogonal planes and comparisons with Allen Mouse Brain Atlas reference images. From left to right: Allen sagittal reference image; STITCH sagittal section (X = 3.18), clearly showing the hippocampal CA fields and the arrow-like DG; STITCH horizontal section (Y = 4.70); STITCH coronal section (Z = 8.60); Allen coronal reference image. **e**, 2D cell annotations of the virtual slice S36.5 and magnified comparisons of the hippocampal region. From left to right: authentic slice S36, isoST-generated S36.5, SpatialZ-generated S36.5, STITCH-generated S36.5, and authentic slice S37. Magnified views (bottom row) demonstrate that existing methods (isoST, SpatialZ) produced anomalous duplicated hippocampal structures, whereas STITCH accurately reconstructed a singular hippocampal structure in the virtual slice S36.5. **f**, Quantitative leave-one-out benchmarking (reconstructing S36) on a small sub-dataset (S34, S35, S36, S37, S39). Top row: Evaluation of transcriptomic fidelity (box plots for cell PCC and gene PCC, and a bar plot for MSE). Middle row: Evaluation of discrete cell annotation consistency (ACC, NMI, ARI). Bottom row: Evaluation of 3D spatial coordinate geometric precision (CD, HD, HD95). SpatialZ was excluded from the geometric evaluation because its generated flat slices cannot match the authentic CCF coordinates. STITCH outperformed SpatialZ and isoST across all metrics. **g**, Comparison of training and inference computation time. STITCH demonstrated superior efficiency, requiring only 0.44 hours for training on the context slices (compared to 2.55 hours for isoST) and 3.18 seconds in total for the target slice inference (1.56 seconds for a single-pass Gene Flow, and 1.62 seconds for Structure Flow).

To verify the anatomical continuity of this massive 3D atlas, we performed digital sectioning across three orthogonal planes and compared them with reference images from the Allen Brain Atlas (Fig. 4d). In both the sagittal (X = 3.18) and coronal (Z = 8.60) planes, STITCH reconstructed the complex brain structures with high fidelity. Notably, in the hippocampal region, STITCH accurately captured the anatomical characteristics from different perspectives, reconstructing the clear arrow-like structure of the dentate gyrus (DG). Both its sagittal and coronal cross-sections matched the corresponding Allen Mouse Brain Atlas reference images with a high degree of fidelity. The continuity in the horizontal plane (Y = 4.70) further confirmed that the 3D spatial structures generated by STITCH consistently align with authentic biological topologies across all dimensions.

To rigorously benchmark generative performance, given the high computational cost of existing spatial generative algorithms (such as isoST [25] and standard SpatialZ) on large-scale data, we extracted a small sub-dataset (comprising slices S34, S35, S36, S37, and S39) from the MERFISH mouse brain dataset. We employed the official fast version of SpatialZ to balance testing efficiency without compromising its cell annotation output [24]. During cross-slice morphological evolution, isoST and SpatialZ can produce topological artifacts in certain tissue structures. To visually demonstrate this, we compared the generation of the virtual slice S36.5, bridging slices S36 and S37 (Fig. 4e). Magnified views clearly reveal that both isoST and SpatialZ produced anomalous “double hippocampus” (i.e., presenting two overlapping CA field structures and misaligned DG regions). In contrast, empowered by STITCH’s capability to capture both geometric contours and gene expression evolutionary rules, the generated S36.5 slice presented an intact, singular hippocampal structure, with the sizes of its DG and CA fields situated in an intermediate transition state between S36 and S37.

Using the same sub-dataset, we next performed a leave-one-out evaluation by masking S36 as the validation target and using S34, S35, S37, and S39 as context slices for model training. As shown in Fig. 4f, evaluated in the raw gene expression space, STITCH comprehensively surpassed both SpatialZ and isoST in continuous transcriptomic fidelity (Cell PCC 25th/50th/75th percentiles reached 0.217/0.382/0.564; Gene PCC reached 0.042/0.107/0.223; MSE dropped to 0.188) and discrete biological identity preservation (ACC 0.345, NMI 0.140, ARI 0.090). Regarding 3D spatial geometric precision — a metric for which SpatialZ was excluded because its virtual slices are restricted to flat XY planes that do not adapt to CCF coordinates (see Supplementary Fig. S2) — STITCH substantially surpassed isoST across the CD, HD, and HD95 metrics. Finally, in terms of overall computational efficiency (Fig. 4g), STITCH demonstrated a significant advantage in both training and inference phases. Training the models on the four context slices required only 0.44 hours for STITCH, compared to 2.55 hours for isoST. Furthermore, the inference generation of the target slice (S36) by STITCH required only 3.18 seconds in total (1.56 seconds for a single-pass Gene Flow, and 1.62 seconds for Structure Flow). This is faster than both isoST (3.97 seconds) and the SpatialZ fast version (28.41 seconds), establishing STITCH’s advantage in large-scale spatiotemporal computational tasks.

### 2.5 STITCH achieves 3D reconstruction across diverse sequencing platforms and supports 2D tissue repair

To further validate the robustness of STITCH across diverse sequencing platforms and its capacity to resolve complex spatial defects, we evaluated our framework on spot-based Visium datasets (human BRCA and DLPFC) [36, 37] and a high-resolution 2D Xenium dataset (mouse brain) [38].

First, on the 3D BRCA dataset (Fig. 5a) [36], standard coordinate interpolation often disrupts the strict spatial grid topology inherent to Visium chips. As demonstrated, the virtual slice generated by SpatialZ exhibited irregular spatial “gaps” and failed to maintain the raw array layout. Empowered by the adaptive modality calibration engine within our 3D Structure Flow, STITCH generated a virtual slice that preserved the regular Visium grid topology. Visually, STITCH accurately reconstructed the spatial compartments of complex microenvironments, including Tumor cells, Fibroblasts, Epithelial cells, and T cells. Concurrently, STITCH faithfully preserved the authentic spatial expression patterns of key marker genes, such as *DCN* [39, 40] (Fig. 5b).

**Figure 5.**
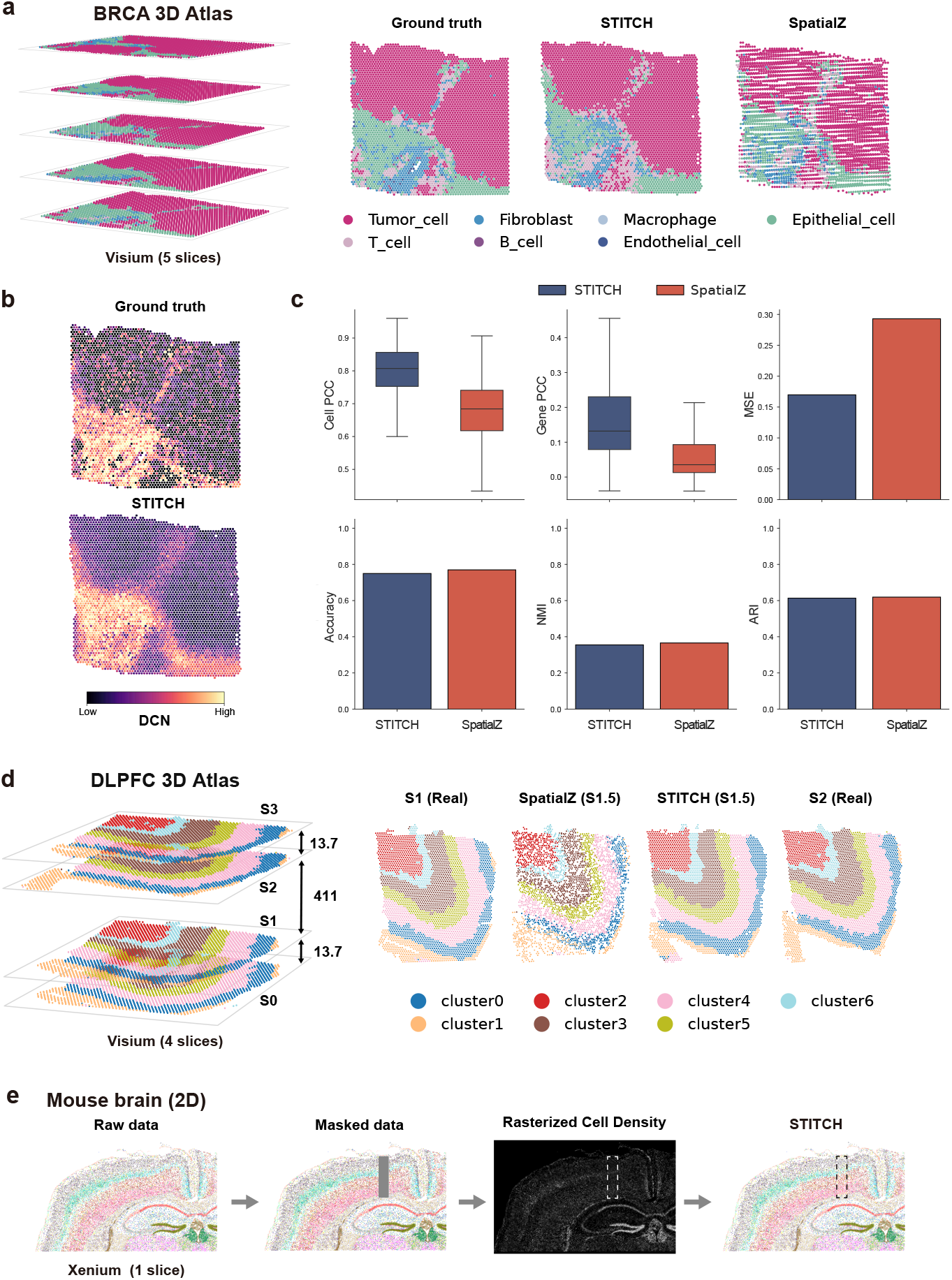
STITCH achieves 3D reconstruction across diverse sequencing platforms and supports 2D tissue repair. **a**, 3D visualization of the Visium BRCA dataset. From left to right: The overall 3D tumor architecture; a selected Ground Truth (GT) slice; the virtual slice reconstructed by STITCH; and the virtual slice generated by SpatialZ. STITCH uniquely maintains the regular grid-array pattern of Visium chips and accurately delineates spatial domains (e.g., Tumor cells, Fibroblasts). **b**, Spatial visualization of the marker gene *DCN*, demonstrating STITCH’s ability to preserve authentic spatial expression patterns. **c**, Quantitative leave-one-out benchmarking between STITCH and SpatialZ in the raw gene space. Boxplots of cell PCC and gene PCC, alongside a bar plot for MSE, highlight STITCH’s significant advantage in transcriptomic fidelity. Categorical metrics (ACC, NMI, ARI) are also provided. **d**, Evaluation on the Visium DLPFC dataset bridging a large spatial gap. The STITCH-generated virtual S1.5 slice preserves the grid layout and accurately reconstructs the tightly packed six cortical layers, whereas SpatialZ produces irregular spatial gaps. **e**, 2D single-slice damage repair on the high-resolution Xenium mouse brain dataset. From left to right: The GT cell annotations; an artificially masked rectangular hole across the cortex; the rasterized cell density image reconstructed through the internal learning strategy; and the final STITCH-reconstructed cell annotations, which reproduce the original cell annotations.

To quantify this transcriptomic fidelity, we conducted a leave-one-out evaluation in the raw gene expression space (Fig. 5c). While SpatialZ held a negligible numerical edge in categorical metrics (e.g., ACC of 0.770 vs. STITCH’s 0.750), STITCH demonstrated significant superiority in modeling continuous gene expressions. STITCH substantially reduced the MSE (0.170 vs. SpatialZ’s 0.293). In terms of gene PCC, STITCH consistently outperformed the baseline across the entire distribution: its 25th, 50th (median), and 75th percentiles reached 0.079, 0.132, and 0.230, respecttively, compared to a mere 0.013, 0.035, and 0.093 for SpatialZ. Cell PCC exhibited a similarly consistent lead (median 0.807 vs. 0.684). These quantitative findings indicate that STITCH successfully captures transcriptomic distribution patterns within the underlying continuous transcriptomic space.

This architectural advantage was further corroborated on the DLPFC dataset [37], which features a wide spatial gap between slices S1 and S2 (Fig. 5d). Bridging this massive fault, the STITCH-generated virtual S1.5 slice successfully preserved the regular grid-array pattern, in contrast to the irregular voids observed in SpatialZ, and clearly reconstructed the tightly organized anatomical arrangement of the six cortical layers.

Beyond 3D cross-slice generation, repairing tissue damage within a single 2D slice serves as a critical scenario to validate the framework’s versatility. To rigorously benchmark this capability against an intact ground truth, we utilized the high-resolution Xenium 2D mouse brain dataset (Fig. 5e, left panel) [38]. We synthetically masked a rectangular region across the cortex to artificially create an intra-slice gap (Fig. 5e, second panel), closely mimicking the severe physical tissue tears or detachment that occur during actual experimental sectioning. Driven by an internal learning strategy [28], our attention-enhanced diffusion model accurately captured the texture and density priors from the surrounding intact context. As illustrated by the generated rasterized cell density image (Fig. 5e, third panel), the diffusion model effectively restored the subtle density distribution differences across various cortical layers. The final cell annotations, generated based on these diffusion-inferred coordinates (Fig. 5e, right panel), exhibited strong topological concordance with the original undamaged ground truth. These findings not only establish STITCH as a highly versatile multidimensional reconstruction tool operating without external reference atlases, but also further demonstrate the flexibility of STITCH’s decoupled architecture: its core Gene Flow generation module can accommodate disparate spatial coordinate models tailored for multi-dimensional spatial gaps.

## 3 Discussion

Spatial transcriptomics (ST) mapping is limited by physical sectioning and in-slice tissue damage, inevitably leaving massive inter-slice gaps that obscure the continuous spatial organization of biological tissues. In this study, we introduced STITCH, a scalable and robust generative framework designed to reconstruct multi-dimensional virtual ST data. Operating effectively in the absence of external reference atlases or high-resolution histological images, STITCH models intrinsic spatial-transcriptomic distributions within the tissue context. Crucially, driven by a point-wise conditional flow matching model, STITCH achieves strong computational scalability, expanding a sparse MERFISH mouse brain dataset into an high-resolution, continuous 3D atlas comprising over 11.5 million cells in approximately 5 hours on a single commodity GPU.

The robust performance of STITCH is largely attributed to its decoupled architectural design. By explicitly separating spatial structure restoration from gene expression generation, STITCH enables flexible integration of heterogeneous reconstruction tasks. This decoupling allows the framework to integrate disparate dimensional tasks, namely bridging 3D cross-slice gaps via optimal transport-conditioned Structure Flow and repairing 2D in-slice structural damage via an internal learning strategy, while using a unified Gene Flow module for transcriptomic generation across reconstructed regions. Furthermore, synchronizing the evolution of spatial morphology with transcriptomic spatial distributions remains challenging during cross-slice inference. This difficulty becomes particularly evident in anatomically curved structures represented in the Allen CCF space. Our results demonstrate that STITCH effectively alleviates this challenge and reduces topological artifacts such as the “double hippocampus”. This performance arises from the synergistic interaction between the Structure Flow and Gene Flow modules: the Structure Flow reconstructs coherent spatial structure, while the Gene Flow preserves transcriptomic continuity across reconstructed regions, thereby improving consistency between tissue morphology and biological identities.

The reconstruction of the whole-mouse-brain MERFISH dataset further highlights an important practical consideration for million-cell-scale generative modeling: the distribution of training cells does not necessarily reflect the structural complexity of different anatomical regions. While some large brain regions occupy substantial fractions of the dataset and exhibit relatively smooth spatial variation, smaller regions may contain intricate anatomical structures yet comprise only a limited number of cells. The hippocampus provides a representative example. Although it accounts for only a small fraction of the whole-brain dataset, it contains structurally complex and biologically important subregions. We therefore adopted importance sampling to increase the training fraction of hippocampus-associated cells while maintaining whole-brain sampling coverage. This training configuration did not alter the point-wise architecture of STITCH or introduce external atlas priors during inference. The corresponding uniform-sampling ablation showed that the global brain morphology remained continuous, but the boundary between the CA fields and DG became less sharply resolved (Supplementary Fig. S1a,b). These results suggest that, for large-scale ST reconstruction, adjusting the training fraction according to regional structural complexity can help preserve both global continuity and local anatomical fidelity.

To develop STITCH, we further evaluated the influence of local spatial neighborhoods on generative dynamics. Comparative experiments with the neighborhood-enhanced variant (STITCH-n) showed that incorporating local neighborhood information provides moderate improvements in reconstruction quality, but substantially increases computational and memory costs. To balance reconstruction accuracy and computational efficiency, we ultimately adopted the decoupled point-wise STITCH architecture as the default framework. This design achieves linear computational complexity while avoiding the severe memory overhead associated with large-scale spatial graph construction, thereby maintaining competitive reconstruction performance with substantially improved scalability.

Despite its robust capabilities, STITCH has specific inherent limitations. The framework implicitly assumes that neighboring intact regions contain sufficient information to support reconstruction of missing areas. Consequently, as observed in our progressively masked validations, generation performance inevitably decays as the physical 3D gaps become excessively large. If a missing spatial span is wide enough to encompass an entirely distinct, highly localized anatomical sub-region that leaves no trace in the adjacent reference slices, STITCH cannot synthesize this unobserved biological structure out of nothing. This limitation, however, is not specific to STITCH but applies universally to all reconstruction methods.

Looking forward, as spatial biology rapidly evolves, the scalable and modular nature of STITCH offers broad prospects for future extensions. The flow matching framework proposed here is not limited to spatial transcriptomics; it can potentially be generalized to broader spatial multi-omics modalities, such as spatial proteomics. Additionally, as an open generative foundation, STITCH can be flexibly integrated with multi-modal priors (such as high-resolution histological images) in the future to achieve even more accurate generation. Overall, STITCH provides a scalable cross-platform framework for reconstructing continuous spatial transcriptomic atlases from sparse and discontinuous tissue observations.

## 4 Methods

### 4.1 Overview of STITCH framework

STITCH is a decoupled generative framework specifically designed for spatial transcriptomic virtual data reconstruction. To bypass the computational bottlenecks of modeling high-dimensional features and spatial coordinates simultaneously, the framework is architected into a three-stage sequential pipeline.

#### Encoder-Decoder module

By compressing high-dimensional gene expressions into a low-dimensional latent representation using a spatial-aware graph autoencoder (SAGA), this module preserves local spatial topologies while deeply integrating spatial coordinates with transcriptomic profiles.

#### Structure Flow module

Operating in the physical space, this module adaptively reconstructs missing morphologies for both 3D cross-slice gaps and 2D single-slice damages. It employs optimal transport-conditioned flow matching to bridge 3D cross-slice gaps, and utilizes an internal learning trategy to repair 2D single-slice damages.

#### Gene Flow module

Conditioned on the reconstructed spatial geometry, this module generates the corresponding transcriptomic profiles in the latent space through point-wise conditional flow matching.

Through this decoupled design, STITCH separates spatial structure reconstruction from transcriptomic generation, improving computational scalability for large-scale spatial reconstruction.

### 4.2 Encoder-Decoder Module

Let 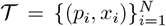 represent a spatial transcriptomic tissue containing *N* cells or spots, where *p*_*i*_ ∈ ℝ^*d*^(*d* ∈ {2, 3}) is the spatial coordinate and *x*_*i*_ ∈ ℝ^*m*^ is the corresponding high-dimensional gene expression profile. We designed an Encoder-Decoder module to reduce transcriptomic dimensionality while preserving multidimensional spatial topologies based on a spatial-aware graph autoencoder (SAGA) [41]. In SAGA, we first constructed a spatial connectivity graph based on the spatial coordinates {*p*_*i*_}. The spatial neighborhood 𝒩_*i*_ of a target node *i* is defined using both a radius-based threshold *r* and a *K*-nearest neighbors (KNN) strategy:

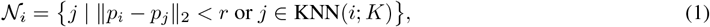

where ∥ · ∥_2_ is the length of a vector. Building on this spatial graph, SAGA employs a multi-layer Graph Attention Network (GAT) as an encoder to compress the high-dimensional transcriptomic features {*x*_*i*_} into low-dimensional latent representations {*z*_*i*_}, which are then used by a decoder to produce reconstructed features 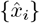.

#### Encoder

Formally, let 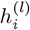 denote the hidden state of node *i* at layer *l*, with 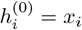 being the initial transcriptomic features. At the *l*th layer of GAT [42], SAGA explicitly encodes spatial distances and directional vectors into spatial edge features. For a target node *i* and its neighbor *j* ∈ 𝒩_*i*_, we concatenated their relative spatial coordinate difference Δ*p*_*ij*_ = *p*_*i*_ − *p*_*j*_ and the distance *d*_*ij*_ = ∥Δ*p*_*ij*_∥_2_, which are then passed through a multi-layer perceptron (MLP) to generate the spatial feature representation:

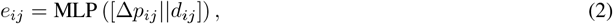

where [·||·] denotes vector concatenation.

To account for the anisotropic spatial structure commonly observed in 3D ST datasets, we employed decoupled intraslice and inter-slice distance decay parameters to explicitly account for the substantial differences between intra-slice (XY-plane) resolution and inter-slice (Z-axis) sectioning thickness by setting:

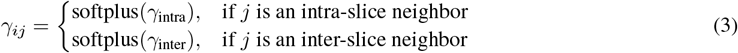

where *γ*_intra_ and *γ*_inter_ are learnable parameters guiding the anisotropic message passing. We further incorporated spatial distance constraints into the attention computation to preserve local spatial consistency. Given the node hidden representations 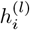 and 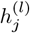 at layer *l*, alongside the spatial edge feature *e*_*ij*_, the preliminary unnormalized attention score is computed as:

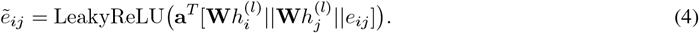

And then, SAGA explicitly computes the attention weights using decoupled anisotropic decay parameters *γ*_*ij*_, which promotes local spatial consistency during message passing:

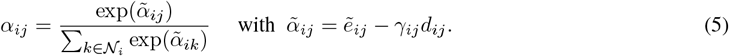

The latent representations are then aggregated and updated via this spatial-aware attention mechanism:

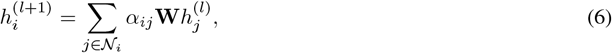

where **W** denotes the learnable feature transformation matrix. For clarity, the above formulation presents a single attention head. In practice, SAGA employs multi-head attention in intermediate encoder layers. The output of the *m*th attention head is computed as

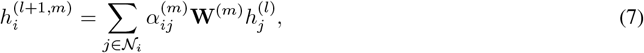

where 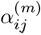 and **W**^(*m*)^denote the attention coefficient and feature transformation matrix associated with the *m*th attention head, respectively. The outputs of all attention heads are subsequently concatenated and transformed through a nonlinear activation:

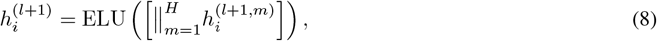

where *H* denotes the number of attention heads, and ELU represents the ELU activation. The final encoder layer adopts a single attention head without feature concatenation to generate the latent representation.

#### Decoder

The latent representation generated by SAGA is subsequently mapped back to the original gene expression space through an MLP decoder. The decoder consists of a series of fully connected layers with ELU activations and dropout regularization. Given the latent embedding {*z*_*i*_}, the reconstructed expression profile is obtained as

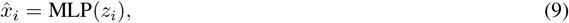

where MLP(·) denotes the decoder network and 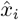 represents the reconstructed gene expression vector.

#### Loss Function

To preserve transcriptomic fidelity and latent manifold structure across diverse spatial resolutions, SAGA is optimized using a composite objective:

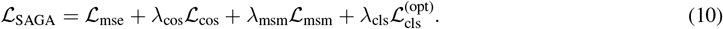

The objective combines reconstruction fidelity (ℒ_mse_ and ℒ_cos_), global manifold preservation (ℒ_msm_), and optional annotation-guided regularization 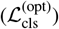.

1. **Reconstruction Losses (ℒ**_**MSE**_ **and ℒ**_**cos**_). The mean squared error preserves absolute gene expression magnitudes, while the cosine loss maintains transcriptomic directional consistency:

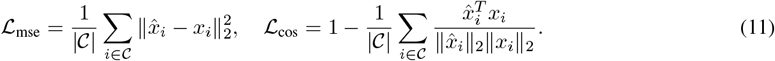
2. **Manifold Similarity Matching Loss (ℒ**_**msm**_). To preserve global transcriptomic topology, we align pairwise cosine similarity matrices between the original expression space and the latent space:

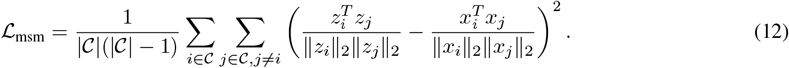
3. **Annotation-Guided Classification Loss (ℒ**_**cls**_). For spot-based platforms, an auxiliary classification objective is optionally applied to enhance latent category separability:

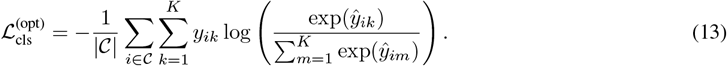

where *ŷ*_*ik*_ denotes the predicted logit for category *k*, and *y*_*ik*_ is the corresponding ground-truth indicator.

For large-scale atlas reconstruction, mini-batch subgraph sampling is employed to prevent memory overflow, and all loss terms are computed exclusively on center nodes with complete local neighborhoods to avoid graph truncation artifacts.

### 4.3 Structure Flow Module

Before reconstructing transcriptomic features, STITCH must first infer the spatial morphology of the missing or damaged tissue regions. To accommodate 3D reconstruction and 2D repair tasks in spatial transcriptomics data, we designed two independent coordinate generation schemes operating within the physical space.

#### 4.3.1 3D Cross-Slice Coordinates Generation

Given two adjacent authentic observed slices denoted as *S*_prev_ and *S*_next_, we modeled the cross-slice spatial morphological transition as a continuous conditional flow matching problem [30].

##### FGW-OT Matching

To establish cross-slice coordinate correspondences, STITCH uses fused Gromov-Wasserstein (FGW) optimal transport [43, 44], which jointly considers absolute spatial displacement and intra-slice spatial topology. Compared with purely distance-based matching, FGW-OT better preserves relative tissue geometry during morphological evolution and reduces mismatches caused by local geometric ambiguity.

For the source slice 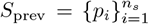 and the target slice 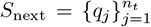, we define the inter-slice physical cost matrix 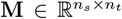 based on normalized squared Euclidean distance. To capture intra-slice spatial topology, we construct a KNN spatial graph for each slice and compute the geodesic distance matrices 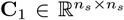 and 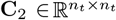 via the shortest path algorithm. STITCH then solves the fused optimal transport problem:

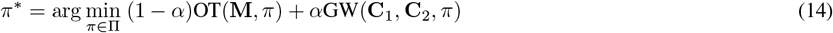

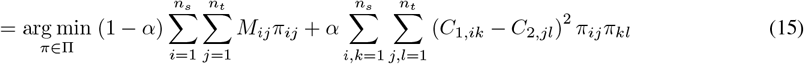

for the transport plan 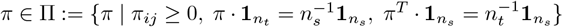, where *α* ∈ [0, 1] controls the trade-off between absolute physical alignment and relative structural preservation. Matched coordinate pairs are then derived from the optimized transport plan *π*^∗^for subsequent Structure Flow training. To eliminate spurious outliers, we implement a robust statistical filter on the matched pairs (Supplementary Note 1).

For large-scale datasets, STITCH uses representative landmark subsets to reduce the computational cost of FGW-OT. For anatomically complex datasets, biological annotations can be further introduced as optional structural priors to reduce cross-boundary mismatches. Under this setting, landmarks are extracted according to cell annotations, and FGW-OT matching is performed within corresponding annotation groups. In the MERFISH mouse brain dataset, STITCH combined these two strategies by performing cell-annotation-guided landmark FGW-OT matching.

##### Continuous Structure Flow Matching

Based on the matched coordinate pairs (*p*^(0)^, *p*^(1)^) ~ *π*^∗^, we define a deterministic linear interpolation path for *t* ∈ [0, 1]. The conditional transport path of the spatial coordinates is formulated as:

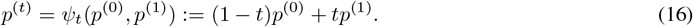

The corresponding conditional structure vector field is defined as the time derivative of the interpolation path:

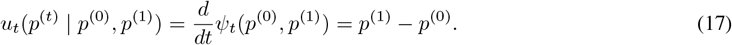

To parameterize and fit this vector field, we construct a neural network *u*_*θ*_(*t, γ*(*p*^(*t*)^)), where the instantaneous spatial coordinates are first mapped via a Fourier positional embedding *γ*(*p*^(*t*)^) to better represent high-frequency spatial variations:

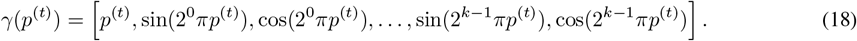

We employed adaptive layer norm (adaLN) blocks [45, 46], where the temporal variable *t* is transformed into adaptive gate (*α*), shift (*β*), and scale (*γ*) parameters to modulate the hidden representations:

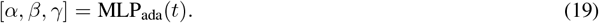

In each adaLN block, the hidden spatial representation *h* is modulated and processed through a feed-forward network (FFN):

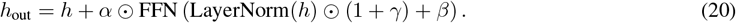

After stacked adaLN blocks, the final hidden representation *h*_final_ is projected to the spatial velocity field:

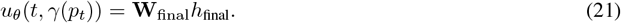

Following the adaLN-Zero initialization strategy [46], the network parameters are optimized by minimizing the mean squared error against the conditional vector field:

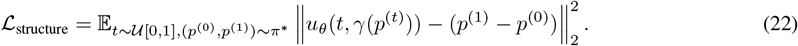

##### Bidirectional ODE Inference

During inference, STITCH performs bidirectional ordinary differential equation (ODE) integration [47] from both neighboring slices. Forward integration is initialized from *S*_prev_ (*t* = 0 → *t*_target_), while backward integration is initialized from *S*_next_ (*t* = 1 → *t*_target_), producing two independently inferred coordinate sets:

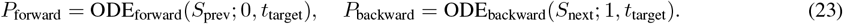

After outlier removal, the final virtual slice is constructed by proportionally sampling and merging points from the forward and backward inferred coordinate sets according to the target interpolation ratio *r* = *t*_target_ ∈ [0, 1], followed by random downsampling to the target cell count.

Finally, platform-specific modality calibrations (e.g., Statistical Outlier Removal for single-cell platforms or Dynamic Grid Snapping and Calibration for array-based data) applied to obtain the calibrated coordinates are in Supplementary Note 1.

#### 4.3.2 2D In-Plane Coordinates Generation

For structural voids or low-quality damaged regions within a single 2D slice, cross-slice interpolation frameworks fail due to the absence of adjacent morphological constraints. To address this problem, STITCH reformulates the discrete coordinate restoration task into an internal image inpainting problem. Building upon the SinDiffusion [28], we extended this framework to an attention-enhanced diffusion model.

##### Cell Density Rasterization

To bridge discrete coordinate point clouds with image-based generative architectures, we first project the 2D spatial positions of the undamaged tissue context into a rasterized single-channel cell density map *I* ∈ ℝ^*H*×*W*×1^. The intensity value of each pixel (*i, j*) represents the local cell density:

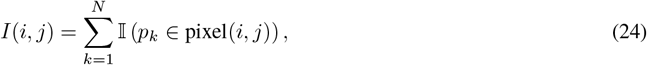

where 𝕀(·) is the indicator function.

##### Attention-Enhanced Diffusion on Density Map

Let the original undamaged density map be denoted as *y*_0_ ~ *q*(*y*_0_). We established a forward diffusion process that gradually adds Gaussian noise over *T* steps according to a variance schedule *β*_*t*_:

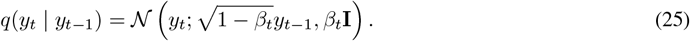

The generative reverse process is parameterized by neural networks *µ*_*θ*_(*y*_*t*_, *t*) and Σ_*θ*_(*y*_*t*_, *t*) to approximate the true posterior, recovering the density map from pure noise *x*_*T*_ :

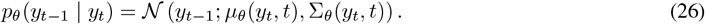

While we adhere to the SinDiffusion framework, the original training mechanism must be restructured to accommodate the fractured nature of spatial transcriptomics data. Specifically, the original SinDiffusion is limited to training on a single, complete image. Applying it directly to slices with missing parts would inevitably introduce invalid void pixels into the training pool, thereby corrupting the generative distribution. To address this issue, we redesign the sampling strategy such that the network is optimized exclusively on patches randomly sampled from valid tissue regions of *I*, ensuring a learning of the sample-specific distribution. Furthermore, we additionally incorporated self-attention layers into the U-Net backbone of SinDiffusion to better capture long-range spatial dependencies.

##### Masked Inpainting Inference

During the inference phase, the undamaged context surrounding the tissue void serves as a spatial boundary constraints to guide the reverse generation. Let *M* ∈ {0, 1} ^*H*×*W*^ be the binary damage mask, where *M* = 1 indicates the authentic intact region and *M* = 0 designates the missing hole. At each reverse denoising step *t*, we enforce a masked replacement strategy:

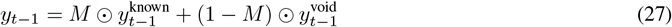

where 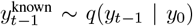 and 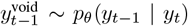. By replacing the known regions with appropriately noised ground-truth observations at each denoising step, the model is compelled to synthesize density patterns exclusively for the missing region that remain consistent with the surrounding spatial context, ultimately yielding the completed density map *Î*_void_. The completed density map is subsequently transformed back into cell-level spatial coordinates using a low-discrepancy quasi-Monte Carlo (QMC) decoding strategy (Supplementary Note 2).

### 4.4 Gene Flow Module

After reconstructing spatial coordinates *P*_target_, STITCH reconstructs transcriptomic latent features through a conditional flow matching framework [30].

Let 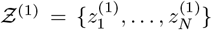 denote the target latent distribution encoded by SAGA (corresponding to *t* = 1). To construct the prior distribution *Ƶ* ^(0)^, we randomly sample a contextual subset *G* ⊂ {1, 2, …, *N*}, and partition the spatial nodes into a contextual reference set *G* (authentic, undamaged cells) and a target generation set *G*^*c*^(missing cells). The reference set *G* serves as the stationary boundary condition of the conditional generative flow, meaning its initial latent state is fixed to its terminal state: 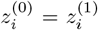 for *i* ∈ *G*. Meanwhile, the target generation region *G*^*c*^is initialized from standard Gaussian noise: 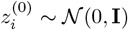 for *i* ∈ *G*^*c*^.

For each individual node *i* ∈ *G*^*c*^, the conditional transport path of the latent state is formally defined as a deterministic linear interpolation:

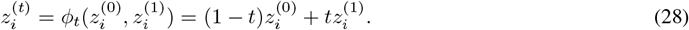

According to conditional flow matching theory, the conditional latent vector field is defined as the time-derivative of this path:

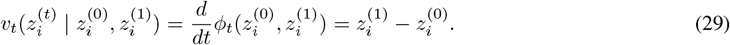

Under this formulation, the target velocity for the stationary boundary nodes (*i* ∈ *G*) remains zero (i.e., *v*_*t*_ = 0).

#### 4.4.1 STITCH: Scalable Point-wise Conditional Flow Matching

To enable scalable reconstruction of large-scale atlases, STITCH models transcriptomic reconstruction through fully decoupled node-wise latent trajectories. In this formulation, the latent velocity field of each node is parameterized conditioned only on its own latent state, temporal condition, and spatial coordinate:

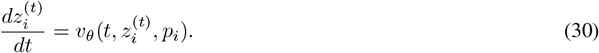

Architecturally, we define a spatiotemporal condition signal *s*_st_ = MLP_time_(*t*) + MLP_space_(*p*_*i*_). The latent state 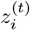 is first embedded into a hidden representation *h*. The velocity network is then constructed using stacked adaLN blocks. For each adaLN block, *s*_st_ is projected into gate (*α*), shift (*β*), and scale (*γ*) parameters:

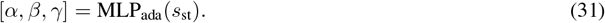

The hidden representation is modulated and processed through a FFN like (20). After stacked adaLN blocks, the final hidden representation *h*_final_ is projected to the velocity field:

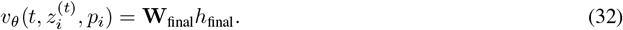

The corresponding point-to-point flow matching optimization objective is defined as:

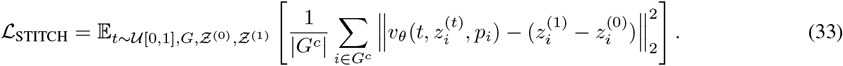

Through this fully decoupled node-wise design, STITCH achieves linear computational complexity.

#### 4.4.2 STITCH-n: Neighborhood-Coupled Conditional Flow Matching

While the independent point-wise generation of STITCH achieves high scalability, explicitly modeling local spatial microenvironments offers a direct mechanism to capture fine-grained transcriptomic gradients and local contextual dependencies. To this end, we developed a neighborhood-enhanced variant, STITCH-n, which explicitly incorporates coupled spatial interactions into the conditional flow dynamics.

In STITCH-n, nodes are no longer conditionally independent. Instead, the velocity field of a target node *i* ∈ *G*^*c*^is explicitly modeled as a coupled function of its own state, its spatial coordinate *p*_*i*_, and the instantaneous latent states of its neighborhood. This neighborhood is naturally partitioned into two subsets: the dynamically evolving target neighbors and the stationary authentic contextual neighbors. The velocity field is therefore formulated as:

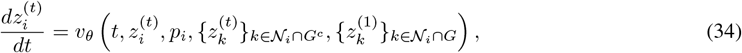

where 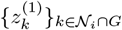 represents the fixed, authentic latent representations of the reference nodes serving as stationary boundary constraints.

Since neighborhood interactions are jointly modeled during flow evolution, all nodes must share a unified global temporal variable *t*. To model these neighborhood interactions, we compute a neighborhood context vector *c*_*i*_ via a Time-Aware Context Attention mechanism. Let *γ*(*t*) denote the temporal embedding. The query, key, and value vectors are computed as:

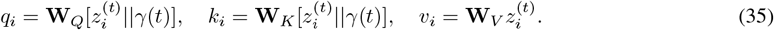

Note that for stationary neighbors (*i* ∈ *G*), their latent state satisfies 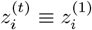 across all time steps. The normalized attention coefficients are computed over a sparse top-*K* neighborhood:

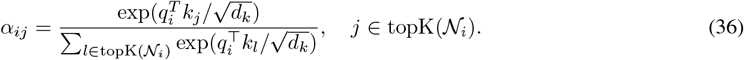

where *d*_*k*_ is the dimension of *q*_*i*_ and *k*_*j*_, and topK(𝒩_*i*_) denotes the subset of *K* neighbors in 𝒩_*i*_ with the largest scaled attention scores 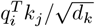. The above equation is written in a single-head form; in practice, STITCH-n implements this operation using multi-head attention. The aggregated neighborhood feature is then computed as 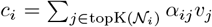. This neighborhood representation and the baseline spatiotemporal condition are then fused in the parameter space through decoupled adaLN conditioning:

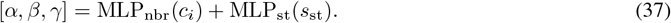

The velocity field is then constructed using the same adaLN-based velocity architecture as in STITCH. The neighborhood-coupled flow matching loss function for STITCH-n is defined as:

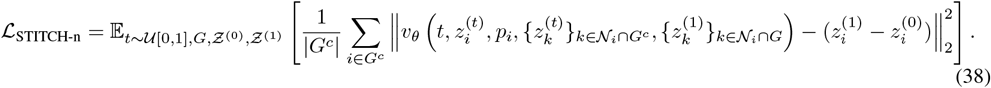

### 4.5 Evaluation Metrics

Because the virtual spatial coordinates generated by diverse models form distinct point clouds that do not overlap with the spatial ground truth (GT) coordinates, we employed a 1-Nearest Neighbor (1-NN) projection strategy to ensure a rigorous cell-to-cell comparative evaluation. Specifically, the generated virtual dataincluding both continuous gene expression profiles and discrete cell type labelswere projected onto the exact spatial coordinates of the GT reference using distance-weighted 1-NN mapping. All subsequent quantitative metrics were calculated between these spatially aligned predictions and the GT references.

#### Transcriptomic Fidelity Metrics

To evaluate the capability of continuous gene expression reconstruction, we utilized three primary metrics: mean squared error (MSE), cell Pearson correlation coefficient (cell PCC), and gene Pearson correlation coefficient (gene PCC).

1. **MSE:** Calculated globally across the entire flattened gene expression matrices to measure the absolute numerical deviation between the aligned generated expressions and the authentic GT expressions.
2. **Cell PCC:** Computed row-wise for individual cells. It measures how well the generated transcriptomic profile of a specific cell correlates with its true biological profile across all genes.
3. **Gene PCC:** Computed column-wise for individual genes. In spatial omics, gene PCC is exceptionally critical as it quantifies the accuracy of reconstructing the macroscopic spatial distribution pattern of a specific gene across the entire tissue architecture.

To maintain numerical stability during PCC calculations, genes or cells with zero variance were dynamically excluded. Detailed subsampling strategies for cell PCC visualization to optimize memory constraints across varied datasets are provided in the Supplementary Information.

#### Biological Identity Metrics

To assess the preservation of discrete biological identities, virtual cells were first assigned cell type labels using a Random Forest classifier [48] trained on the surrounding known context slices. After the aforementioned 1-NN spatial alignment, these predictions were compared against the GT annotations using:

1. **Accuracy (ACC):** The proportion of spatially aligned virtual cells whose predicted cell types match the GT labels.
2. **Adjusted Rand Index (ARI):** A metric evaluating the similarity between the predicted cell type clustering and the true clustering, adjusted for the chance grouping of elements.
3. **Normalized Mutual Information (NMI):** An information-theoretic measure quantifying the mutual dependence between the predicted and GT labels, normalized to scale between 0 and 1.

#### Spatial and Geometric Metrics

To quantify the morphological precision of the reconstructed 3D spatial coordinates, we treated the GT coordinates (*T*) and the generated virtual coordinates (*P*) as two distinct point clouds and calculated geometric distances using KD-Trees:

1. **Chamfer Distance (CD):** The symmetric average of the nearest neighbor distances between the two point clouds, defined as the mean distance from each point in *P* to its closest point in *T*, plus the mean distance from each point in *T* to its closest point in *P*.
2. **Hausdorff Distance (HD):** Measures the maximum geometric mismatch (i.e., the worst-case boundary error) between the two point clouds. It is defined as the maximum of all minimum distances from *P* to *T* and *T* to *P*.
3. **95th Percentile Hausdorff Distance (HD95):** To mitigate the disproportionate impact of extreme outliers on the standard HD, we calculated the 95th percentile of the nearest neighbor distances, providing a more robust evaluation of macroscopic morphological consistency.

The rigorous mathematical definitions and specific dynamic subsampling strategies for all the aforementioned continuous transcriptomic and geometric metrics are comprehensively detailed in Supplementary Note 3.

### 4.6 Datasets and Platform-Specific Preprocessing

To comprehensively evaluate the robustness and generalizability of the STITCH framework across real and diverse research scenarios, we curated five publicly available spatial transcriptomics datasets. These datasets span different sequencing modalities, resolutions, and species. Given the fundamental differences in the underlying data modality between whole-transcriptome sequencing and targeted in situ hybridization (ISH), we avoided a rigid, one-size-fits-all preprocessing pipeline. Instead, we performed platform-specific processing to optimally preserve the intrinsic biological signals of each technology. Furthermore, because the absolute values of spatial coordinates exported from different platforms vary considerably, we applied moderate numerical scaling where necessary to ensure numerical stability during neural network training.

#### Whole-Transcriptome Sequencing Datasets

Array or DNA nanoball-based whole-transcriptome technologies typically capture tens of thousands of genes, accompanied by substantial background noise; therefore, feature selection is a necessary standard step.

#### Visium spot-level datasets (BRCA and DLPFC)

Both are 10x Genomics Visium datasets obtained as 3D spatially aligned reconstructions from the So3D database [49]. The breast cancer (BRCA) data, comprising 21,154 spatial spots across 17 sections from a larger pan-cancer cohort, originally come from Mo et al. [36]. The dorsolateral prefrontal cortex (DLPFC) data, containing 14,243 spatial spots, were originally generated by Maynard et al. [37]. For both datasets, the supplied gene expression matrices were already log1p-normalized. We further selected the top 1,500 spatially variable genes (SVGs) across the dataset as model inputs. To ensure numerical stability during neural network optimization while preserving the authentic physical topological architecture, the spatial coordinates provided by So3D were proportionally scaled rather than retaining their raw absolute values.

#### Stereo-seq *Drosophila* embryo dataset

This 3D dataset was obtained from the public data repository provided by the Spateo spatial omics computational framework [10]. Similar to the processing logic for the Visium data, we extracted the top 1,000 SVGs from the log1p-normalized matrix for the 21,473 cells. To rigorously test the model’s adaptability to real physical scales, the spatial coordinates of this dataset were not scaled in any way and were kept in their original physical units.

#### Targeted Gene Panel Datasets

For image-based targeted technologies, the pre-designed gene panels cover a relatively concise set of core genes.

#### MERFISH mouse brain dataset

This MERFISH mouse brain dataset was originally generated by Zhang et al. [4] and was obtained from the Zhuang-ABCA-2 collection in the Allen Brain Cell Atlas (ABC Atlas). The raw data was provided as a comprehensive tabular database rather than discrete slice files. To ensure optimal data quality for robust 3D reconstruction, we performed rigorous quality control by filtering out several severely damaged or low-quality sections. Subsequently, we systematically extracted the spatial coordinates and the corresponding feature matrices for all retained cells, assembling them into a unified 3D H5AD object comprising 54 high-quality slices (totaling 1,056,520 cells). The exact index list of the selected 54 sections is detailed in Supplementary Note 4. This dataset contains 1,122 targeted genes; we utilized the full expression profile as input without further feature selection. For the spatial coordinates, we used the publicly provided coordinates registered to the Allen Mouse Brain Common Coordinate Framework version 3 (CCFv3) [35]. To facilitate visualization and digital sectioning, we reoriented the CCF coordinates into an anatomical coordinate system, where the X, Y, and Z axes correspond to the left-right, superior-inferior, and anterior-posterior directions, respectively.

#### Xenium 2D mouse brain dataset

This dataset was obtained from the 10x Genomics official platform [38]. It comprises a total of 36,602 cells and contains 248 genes. High-fidelity restoration of single 2D sections is a core capability of the STITCH framework for handling tissue damage. To fully demonstrate the high modularity and flexibility of our decoupled architecture, we adopted a highly targeted feature processing strategy for this task. Specifically, we curated a feature subset of 50 core spatially variable genes (SVGs). Because this core gene panel already possesses very low dimensionality in expression space, we directly guided the core generative engine (Gene Flow) to perform conditional flow matching in the original feature space. This design verifies STITCH’s ability to perform end-to-end generation of a core targeted gene set without prior dimensionality reduction. Regarding spatial coordinates, because the original coordinates of this platform are large in absolute values, we applied standard Z-score normalization to the 2D coordinates to ensure numerical stability during training.

### 4.7 Implementation Details and Baseline Configurations

The STITCH framework was implemented in Python utilizing the PyTorch deep learning library. To ensure full transparency and computational reproducibility, the core neural network architectures, optimization strategies, and the dataset-specific hyperparameter configurations specifically designed for the STITCH framework are thoroughly documented in Supplementary Note 5.

To ensure a strictly fair and objective comparison, all baseline methods evaluated in this study were executed using their official open-source implementations, and their hyperparameter settings adhered to the default configurations recommended in their original publications. Notably, for the MERFISH mouse brain dataset, executing the standard SpatialZ algorithm incurred prohibitive runtime on our infrastructure. Consequently, we employed the official “fast” version of SpatialZ for this specific dataset; as documented by its authors, this accelerated version reduces computational overhead while maintaining the accuracy of cell annotation [24]. For isoST cell annotation, we followed the KNN-classifier strategy described in the original isoST study [25]; because the corresponding classifier parameters were not provided in the official repository, we used 15 neighbors and retained the default settings for all other classifier parameters.

Finally, all computational benchmarking reported in this study including the precise measurement of execution time and peak memory consumption across all evaluated models was conducted on a dedicated standalone workstation equipped with an Intel Core i9-14900KF CPU and a single NVIDIA GeForce RTX 4090 GPU.

## Data Availability

All spatial transcriptomics datasets used in this study are publicly available. The Stereo-seq *Drosophila* embryo dataset was obtained from the Spateo computational framework [10] and can be directly downloaded at https://www.dropbox.com/s/bvstb3en5kc6wui/E7-9h_cellbin_tdr_v2.h5ad?dl=1. The MERFISH mouse brain dataset (specifically the ABCA-2 release) is available from the original publication [4] and can be accessed via the Allen Brain Cell (ABC) Atlas portal at https://alleninstitute.github.io/abc_atlas_access/descriptions/Zhuang-ABCA-2.html. To ensure the reliability of the 3D spatial benchmarks, the 3D aligned versions of the Visium human BRCA and DLPFC datasets were downloaded from the So3D database [49], avail able at https://So3D.bio-database.com/download.jsp. The raw data for the Xenium 2D mouse brain dataset is openly accessible via the 10x Genomics official dataset portal at https://www.10xgenomics.com/datasets/fresh-frozen-mouse-brain-for-xenium-explorer-demo-1-standard [38].

## Code Availability

The source code for the STITCH framework is available on GitHub at https://github.com/WANG-SUI/STITCH.

## Acknowledgements

T.L. acknowledges the support of the National Key R&D Program of China under grant 2021YFA1003301, and the National Science Foundation of China under grant 12288101. We also thank the High-performance Computing Platform of Peking University for providing the computational resources for this research.

## Author Contributions

All authors conceived the project. S.W. designed and implemented the algorithm. S.W. and X.W. performed the data analysis. Q.P. participated in methodological discussions. T.L., S.W., and X.W. interpreted the results and wrote the paper. T.L. supervised the research.

## Supplementary Information

Supplementary Information is available for this paper.

