## Supplementary Information for "STITCH: Spatial Transcriptomics Imputation via Flow Matching with Internal Learning"

### Supplementary Note 1: Spatial Structure Optimization and Modality Calibration

Continuous Normalizing Flows (CNFs) inherently output trajectories in a strictly continuous physical domain. To adeptly handle the structural diversity across different spatial transcriptomics platforms and ensure the topological coherence of the generated vector fields, STITCH introduces pre-filtering and adaptive post-calibration modules.

**Robust Statistical Filtering for Optimal Transport Mappings** During the Prior-Guided Local FGW-OT Matching phase in 3D cross-slice evolution, computing the optimal transport plan occasionally generates anomalous long-distance mappings (i.e., “flying points”) due to structural ambiguities. If left untreated, these extreme coordinate pairs translate into massive target deformation velocities ($v_{true} = p_1 - p_0$), producing explosive gradients that destabilize the continuous vector field training. To eliminate these artifacts, we implement a robust statistical filter based on the Median Absolute Deviation (MAD). For each class-constrained matching subset, we compute the distance distribution of all matched coordinate pairs $(p_0, p_1)$. Matches whose distances exceed a dynamic threshold—defined as the median distance plus a scaled MAD factor—are systematically rejected prior to network optimization. This strict filtration ensures the topological coherence of the subsequent morphological evolution.

**Dynamic Grid Snapping and Calibration for Array-Based Data** For array-based technologies like 10x Visium, generated continuous physical trajectories must be strictly anchored to a predefined hexagonal grid layout. To bridge this continuous-to-discrete modality gap, we designed a dynamic grid snapping engine. First, the inherent spot-to-spot distance $d$ is dynamically extracted from the physical spatial context. Based on the spatial boundaries of the generated ODE trajectories ($P_{fwd}$ and $P_{bwd}$), an ideal virtual hexagonal grid is constructed.

To determine the activation of each virtual grid node $v_i$, the engine evaluates its spatial affinity to the continuous point clouds by computing its proximity to the generated trajectories. Crucially, to prevent artificial over-smoothing and preserve authentic biological topologies (such as physical fissures or dense tumor boundaries), the snapping process incorporates rigorous structural constraints. Virtual nodes are selectively activated based on a combination of strict geometric proximity bounds (e.g., distance within $0.65d$) and contextual tissue density requirements (e.g., local microenvironmental support or matching the overall target spot abundance $N_{target}$). This distance-and-density-guided selection guarantees that the activated spots seamlessly form a continuous biological contour while strictly obeying hardware grid constraints. Finally, to ensure absolute 3D planar flatness, the Z-axis coordinate is deterministically assigned via exact linear temporal interpolation: $Z_{exact} = (1-t)Z_{start} + tZ_{end}$.

**Statistical Outlier Removal (SOR) for Cellular-Resolution Data** Unlike Visium data, subcellular or single-cell resolution technologies (e.g., MERFISH, Stereo-seq) exhibit irregular, dense continuous distributions. For these modalities, bidirectional ODE integration occasionally yields a minuscule fraction of spatial "outliers" (isolated cells) deviating from the main tissue topology. To ensure the compactness and boundary smoothness of the generated 3D tissue, we applied the Statistical Outlier Removal (SOR) algorithm. Specifically, for each generated cell $p_i$, we calculate the mean distance $\bar{d}_i$ to its $k$ nearest neighbors. Let $\mu$ and $\sigma$ denote the global mean and standard deviation of these distances across the entire slice. If $\bar{d}_i > \mu + \alpha \cdot \sigma$ (with $\alpha$ serving as a stringency threshold), the cell is identified as an ODE integration artifact and physically removed, thereby maintaining the morphological purity of high-resolution spatial atlases.

### Supplementary Note 2: Coordinate Decoding via Quasi-Monte Carlo Sampling

Transforming the generated continuous density image $\hat{I}_{\text{void}}$ back into discrete cell-level physical coordinates poses a risk of introducing artificial grid-like clustering and amplifying background generative noise. To surmount this, we developed a deterministic decoding mechanism utilizing a piecewise density calibration and a low-discrepancy Quasi-Monte Carlo (QMC) sampling strategy based on the scrambled Halton sequence.

For each target pixel $(x, y)$, the local density intensity is first translated into a raw expected cell count $c_{x,y}$. To suppress background noise and maintain biological plausibility in sparsely populated regions, we apply a piecewise density thresholding strategy: expected counts below a strict blanking threshold ($t_{\text{blank}}$) are forced to zero to prevent noise instantiation, while sparse values ($t_{\text{blank}} \leq c_{x,y} < 1.0$) are conservatively assigned a single cell. For dense regions, $c_{x,y}$ is quantized into an initial integer count $N_{x,y}^{\text{base}}$ through a probabilistic floor operation to preserve expectation:

$$N_{x,y}^{\text{base}} = \lfloor c_{x,y} \rfloor + \text{Bernoulli}(c_{x,y} - \lfloor c_{x,y} \rfloor) \tag{S1}$$

To ensure a natural topological transition and avoid density abruptness at the repair margins, we apply morphological erosion to the damage mask to identify boundary pixels. For these surgical seams, a decay factor $\lambda$ is applied to the initial count to simulate natural tissue thinning. The final adjusted cell count $N_{x,y}$ is formulated as:

$$N_{x,y} = \begin{cases} \lfloor \lambda N_{x,y}^{\text{base}} \rfloor + \text{Bernoulli}(\lambda N_{x,y}^{\text{base}} - \lfloor \lambda N_{x,y}^{\text{base}} \rfloor), & \text{if } (x, y) \text{ is boundary} \\ N_{x,y}^{\text{base}}, & \text{otherwise} \end{cases} \tag{S2}$$

Subsequently, we map these scalar counts back to the continuous physical space. We generate a 2D scrambled Halton sequence $\mathbf{h}_i \in [0, 1)^2$ of length $N_{x,y}$, which fills the sub-pixel space with optimal uniformity, mimicking the natural physical dispersion of biological cells without grid artifacts. Let $L \in \mathbb{R}^+$ denote the physical side length of each pixel, which represents the spatial resolution of the rasterized grid. Let $C_{x,y} \in \mathbb{R}^2$ be the absolute physical center coordinate of the pixel $(x, y)$. For the $i$-th cell generated within this pixel, its final continuous physical coordinate $p_i \in \mathbb{R}^2$ is rigorously reconstructed as:

$$p_i = C_{x,y} + L \cdot \left( \mathbf{h}_i - \begin{bmatrix} 0.5 \\ 0.5 \end{bmatrix} \right) + \epsilon_i, \quad \epsilon_i \sim \mathcal{N} \left( 0, \left( \frac{S}{20} \right)^2 \mathbf{I} \right) \tag{S3}$$

The isotropic Gaussian spatial jitter $\epsilon_i$ is superimposed to break any remaining mechanical lattice artifacts derived from the rasterized grid, successfully recovering the authentic micro-spatial stochasticity inherent in real tissue architectures.

#### Supplementary Note 3: Comprehensive Definitions of Evaluation Metrics

To rigorously quantify the generative fidelity and morphological consistency of STITCH, we employed a suite of metrics spanning continuous transcriptomic fidelity, discrete biological identity, and macroscopic geometric structure. Let $X$ denote the ground truth (GT) gene expression matrix, $\hat{X}$ denote the generated expression matrix, $Y$ denote the GT cell type labels, and $\hat{Y}$ denote the predicted labels. Let $T$ and $P$ denote the GT and predicted 3D point clouds, respectively.

**1. Mean Squared Error (MSE)** MSE measures the global absolute numerical deviation across the flattened expression matrices (with $N$ cells and $M$ genes):

$$\text{MSE} = \frac{1}{N \times M} \sum_{i=1}^{N} \sum_{j=1}^{M} (X_{ij} - \hat{X}_{ij})^2 \tag{S4}$$

**2. Pearson Correlation Coefficient (PCC)** PCC evaluates the linear correlation between generated and authentic profiles. For vectors $u$ and $v$ (representing either a cell's transcriptome or a gene's spatial distribution), it is defined as:

$$\text{PCC}(u, v) = \frac{\sum (u_i - \bar{u})(v_i - \bar{v})}{\sqrt{\sum (u_i - \bar{u})^2 \sum (v_i - \bar{v})^2}} \tag{S5}$$

*Sub-sampling Strategy for PCC Visualization:* Due to the prohibitive memory overhead of rendering high-density violin plots for tens of thousands of individual cells and genes, we employed a dynamic random subsampling strategy during PCC calculations. Specifically, for the Stereo-seq *Drosophila* embryo datasets, we randomly sampled up to 4,000 cells for Cell PCC and 1,000 genes for Gene PCC. For larger-scale datasets (MERFISH mouse brain and Visium cohorts), the threshold was expanded to a maximum of 10,000 cells for Cell PCC and 4,000 genes for Gene PCC. Genes or cells with zero variance were dynamically excluded to maintain numerical stability.

**3. Accuracy (ACC)** Accuracy calculates the proportion of spatially aligned virtual cells whose predicted cell types strictly match the GT labels:

$$\text{ACC} = \frac{1}{N} \sum_{i=1}^{N} \mathbb{I}(y_i = \hat{y}_i) \tag{S6}$$

where $\mathbb{I}(\cdot)$ is the indicator function.

**4. Adjusted Rand Index (ARI)** ARI evaluates the similarity between the predicted cell type clustering and the true clustering, mathematically adjusted for chance. Let $n_{ij}$ be the number of cells of true class $i$ assigned to predicted class $j$, $a_i$ be the total cells in true class $i$, and $b_j$ be the total cells in predicted class $j$:

$$\text{ARI} = \frac{\sum_{ij} \binom{n_{ij}}{2} - \left[\sum_i \binom{a_i}{2} \sum_j \binom{b_j}{2}\right] / \binom{N}{2}}{\frac{1}{2}\left[\sum_i \binom{a_i}{2} + \sum_j \binom{b_j}{2}\right] - \left[\sum_i \binom{a_i}{2} \sum_j \binom{b_j}{2}\right] / \binom{N}{2}} \tag{S7}$$

**5. Normalized Mutual Information (NMI)** NMI is an information-theoretic measure quantifying the mutual dependence between predicted labels and GT labels, scaled between 0 and 1. Utilizing the entropy $H(\cdot)$ and mutual information $I(\cdot;\cdot)$:

$$\mathrm{NMI}(Y, \hat{Y}) = \frac{2 \cdot I(Y; \hat{Y})}{H(Y) + H(\hat{Y})} \tag{S8}$$

**6. Chamfer Distance (CD)** CD evaluates global spatial proximity by calculating the symmetric mean nearest-neighbor distance between the two point clouds $P$ and $T$:

$$d_{CD}(P, T) = \frac{1}{2}\left(\frac{1}{|P|}\sum_{p\in P}\min_{t\in T}\|p - t\|_2 + \frac{1}{|T|}\sum_{t\in T}\min_{p\in P}\|t - p\|_2\right) \tag{S9}$$

**7. Hausdorff Distance (HD)** The symmetric HD captures the worst-case boundary mismatch between the generated morphology and the authentic tissue contour:

$$d_{HD}(P, T) = \max\left(\max_{p\in P}\min_{t\in T}\|p - t\|_2,\ \max_{t\in T}\min_{p\in P}\|t - p\|_2\right) \tag{S10}$$

**8. 95th Percentile Hausdorff Distance (HD95)** To mitigate the impact of extreme spatial outliers on the standard HD, HD95 extracts the 95th percentile ($P_{95\%}$) of the nearest-neighbor distance distribution:

$$d_{HD95}(P, T) = \max\left(P_{95\%}\left(\min_{t\in T}\|p - t\|_2\right),\ P_{95\%}\left(\min_{p\in P}\|t - p\|_2\right)\right) \tag{S11}$$

### Supplementary Note 4: Detailed Index of the MERFISH Mouse Brain Dataset

To construct the robust organ-scale 3D macroscopic atlas demonstrated in our main text, we utilized the MERFISH mouse brain dataset from Zhang et al. (2023). The raw database contains numerous sections; however, to prevent extreme physical distortions or severe technical artifacts from introducing unresolvable interpolation noise into the optimal transport mappings, we performed a rigorous structural quality control. We explicitly retained 54 high-quality, biologically coherent contiguous sections.

For full transparency and to facilitate precise computational reproducibility of our macroscopic generative tasks, the exact original identifiers of the 54 retained slices are cataloged below in ascending anatomical order:

```
Zhuang-ABCA-2.004, Zhuang-ABCA-2.005, Zhuang-ABCA-2.006,
Zhuang-ABCA-2.007, Zhuang-ABCA-2.008, Zhuang-ABCA-2.009,
Zhuang-ABCA-2.010, Zhuang-ABCA-2.011, Zhuang-ABCA-2.012,
Zhuang-ABCA-2.013, Zhuang-ABCA-2.014, Zhuang-ABCA-2.015,
Zhuang-ABCA-2.016, Zhuang-ABCA-2.017, Zhuang-ABCA-2.018,
Zhuang-ABCA-2.019, Zhuang-ABCA-2.020, Zhuang-ABCA-2.021,
Zhuang-ABCA-2.022, Zhuang-ABCA-2.023, Zhuang-ABCA-2.025,
Zhuang-ABCA-2.026, Zhuang-ABCA-2.027, Zhuang-ABCA-2.028,
Zhuang-ABCA-2.030, Zhuang-ABCA-2.031, Zhuang-ABCA-2.032,
Zhuang-ABCA-2.033, Zhuang-ABCA-2.034, Zhuang-ABCA-2.035,
Zhuang-ABCA-2.036, Zhuang-ABCA-2.037, Zhuang-ABCA-2.039,
Zhuang-ABCA-2.040, Zhuang-ABCA-2.041, Zhuang-ABCA-2.042,
Zhuang-ABCA-2.044, Zhuang-ABCA-2.045, Zhuang-ABCA-2.046,
Zhuang-ABCA-2.047, Zhuang-ABCA-2.048, Zhuang-ABCA-2.049,
Zhuang-ABCA-2.050, Zhuang-ABCA-2.051, Zhuang-ABCA-2.052,
Zhuang-ABCA-2.053, Zhuang-ABCA-2.054, Zhuang-ABCA-2.055,
Zhuang-ABCA-2.056, Zhuang-ABCA-2.057, Zhuang-ABCA-2.058,
Zhuang-ABCA-2.059, Zhuang-ABCA-2.060, Zhuang-ABCA-2.061.
```

Notably, the deliberately absent indices in the sequence (e.g., `Zhuang-ABCA-2.024`, `Zhuang-ABCA-2.029`, `Zhuang-ABCA-2.038`, `Zhuang-ABCA-2.043`) represent the missing physical sections. It is precisely across these wider macroscopic spatial vacuums that the STITCH Structure Flow dynamically elevated the generation depth to 19 virtual slices, ensuring seamless structural transition for complex brain domains.

### Supplementary Note 5: Key Hyperparameter Configurations

STITCH contains three trainable components: the encoder-decoder module for latent representation learning, the Structure Flow module for spatial coordinate reconstruction, and the Gene Flow module for transcriptomic generation. To document the dataset-specific configurations of these components, we summarize the key hyperparameters for the encoder-decoder module, Structure Flow module, and Gene Flow module in Supplementary Tables S1, S2, and S3, respectively.

Table S1: Key encoder-decoder module configurations.

| **Dataset** | **Input Dim** | **Hidden Dims** | **Graph Setting** |
|---|---|---|---|
| Stereo-seq *Drosophila* | 1000 | [128, 128, 128] | radius-based graph† |
| MERFISH Mouse Brain | 1122 | [256, 256, 256] | $k_{\text{intra}} = 5$, $k_{\text{inter}} = 4$ |
| MERFISH Mouse Brain subset | 1122 | [256, 256] | $k_{\text{intra}} = 5$, $k_{\text{inter}} = 6$ |
| Visium BRCA | 1500 | [256, 256] | $k_{\text{intra}} = 6$, $k_{\text{inter}} = 4$ |
| Visium DLPFC | 1500 | [256, 256] | $k_{\text{intra}} = 6$, $k_{\text{inter}} = 4$ |

*Note:* The encoder-decoder module used spatial-aware graph convolution with 4 spatial input channels. †For the Stereo-seq *Drosophila* dataset, `align_spatial` was used for Fig. 3 benchmarking with a radius cutoff of 17.0, whereas `3d_align_spatial` was used for Fig. 2 benchmarking with a radius cutoff of 9.0.

Table S2: Key Structure Flow module hyperparameter configurations.

| **Dataset** | **Frequencies** | **Landmarks** | **Matching Setting** |
|---|---|---|---|
| Stereo-seq *Drosophila* | 5 | 15000 | FGW $\alpha = 0.6$, geo-$k = 5$ |
| MERFISH Mouse Brain | 7 | 15000 | FGW $\alpha = 0.6$, geo-$k = 12$ |
| MERFISH Mouse Brain subset | 6 | 15000 | FGW $\alpha = 0.6$, geo-$k = 12$ |
| Visium BRCA | 6 | full slice | OT-based matching |
| Visium DLPFC | 6 | full slice | OT-based matching |

*Note:* For 3D cross-slice spatial reconstruction, Structure Flow used hidden dimension 256, 4 AdaLN blocks, AdamW optimization with a learning rate of $1 \times 10^{-3}$ and weight decay of $1 \times 10^{-5}$. For the attention-enhanced SinDiffusion model in the Xenium mouse brain experiment, rasterized images were trained with image size 256, learning rate $5 \times 10^{-4}$, 1000 diffusion steps, attention resolution 2, 16 head channels, and batch size 16.

Table S3: Key Gene Flow Module hyperparameter configurations.

| **Dataset** | **LR** | **AdaLN Blocks** | **Latent Dim** |
|---|---|---|---|
| Stereo-seq *Drosophila* | $1 \times 10^{-4}$ | 4 | 32 |
| MERFISH Mouse Brain | $1 \times 10^{-3}$ | 5 | 64 |
| MERFISH Mouse Brain subset | $1 \times 10^{-3}$ | 4 | 48 |
| Visium BRCA | $3 \times 10^{-4}$ | 4 | 48 |
| Visium DLPFC | $5 \times 10^{-4}$ | 4 | 48 |
| Xenium 2D Mouse Brain | $1 \times 10^{-3}$ | 4 | 50* |

*Note:* Gene Flow used a fixed hidden dimension of 256, AdamW optimization with cosine annealing scheduling, and weight decay of $1 \times 10^{-5}$. Inference used midpoint ODE integration with 50 steps, and predictions were averaged over three repeated samplings unless otherwise specified. *For Xenium, Gene Flow was performed directly on the top 50 spatially variable genes in the raw expression space.

#### Supplementary Figures

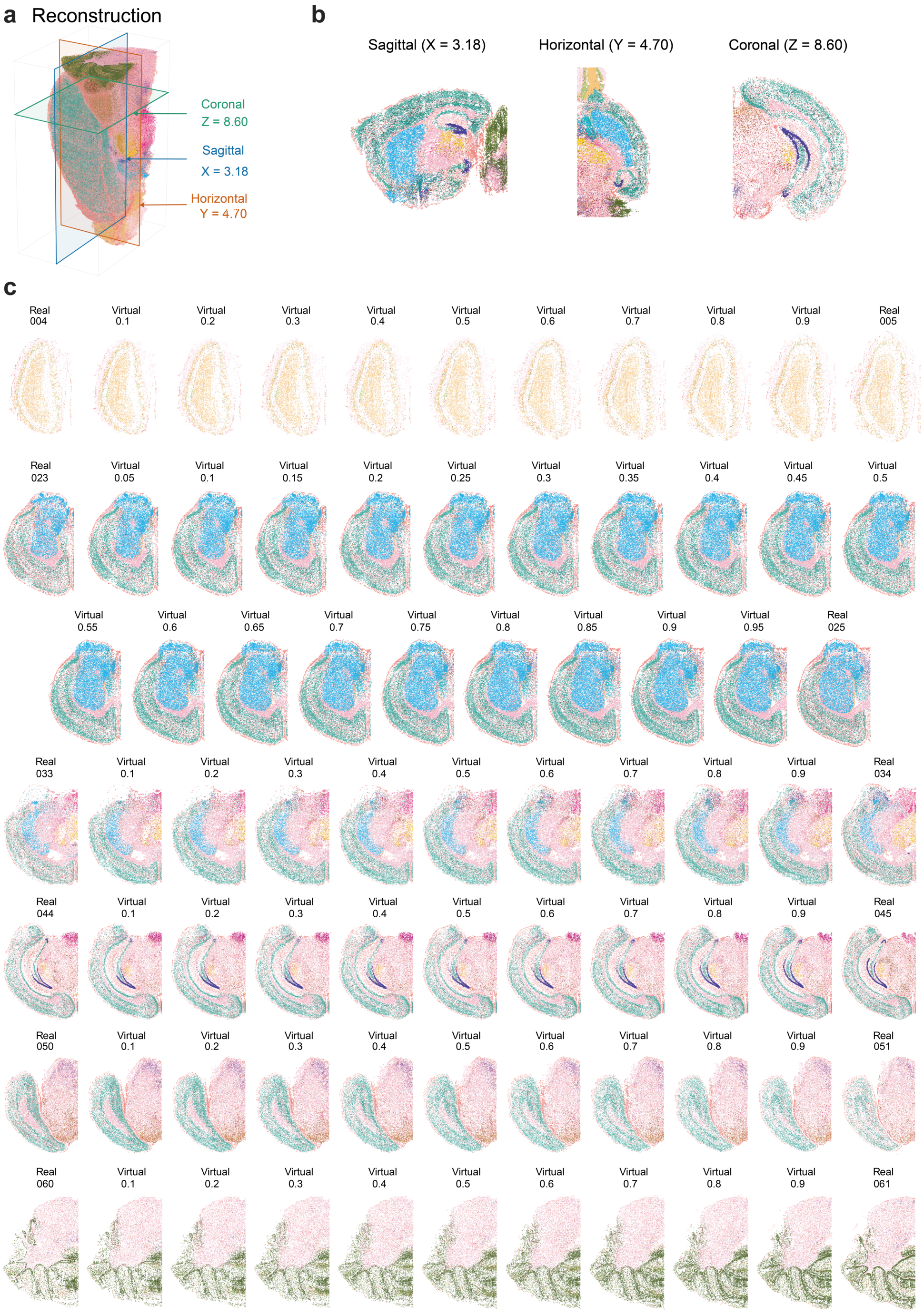


Figure S1: **Supplementary Fig. S1 | Effects of training sampling configuration in whole-brain mouse brain reconstruction. a,** High-resolution 3D continuous mouse brain atlas reconstructed by STITCH using uniform sampling during whole-brain Gene Flow training. **b,** Digital sections of the uniformly sampled reconstruction across three orthogonal planes. **c,** Representative cell annotation maps from selected slice intervals generated by the final whole-brain STITCH configuration, in which the training fraction of hippocampus-associated cells was increased while maintaining whole-brain sampling coverage. The color legend for cell annotations is consistent with that used in **Fig. 4**.

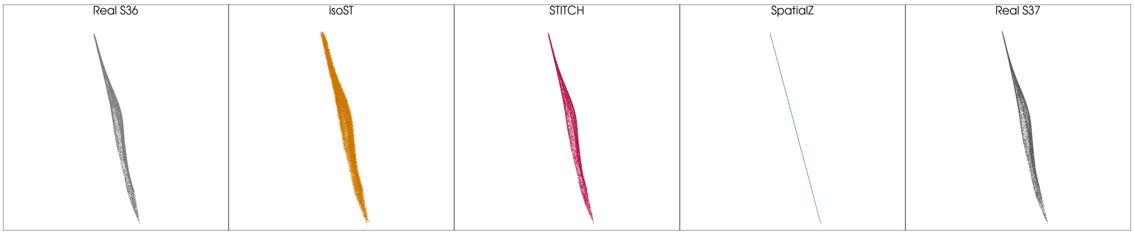


Figure S2: **3D spatial visualizations of authentic and generated slices in the CCF coordinate framework.** The figure displays (from left to right): the authentic slice S36, the isoST-generated S36.5, the STITCH-generated S36.5, the SpatialZ-generated S36.5, and the authentic slice S37.
